# Soluble CTLA-4 mainly produced by Treg cells inhibits type 1 inflammation without hindering type 2 immunity to allow for inflammation resolution

**DOI:** 10.1101/2023.05.26.542386

**Authors:** Motonao Osaki, Shimon Sakaguchi

**Affiliations:** Laboratory of Experimental Immunology, WPI Immunology Frontier Research Center, Osaka University, Suita, Osaka 565-0871, Japan; Laboratory of Experimental Immunology, Institute for Life and Medical Sciences, Kyoto University, Kyoto 606-8507, Japan

**Author notes:** **Corresponding author** Correspondence to Shimon Sakaguchi Laboratory of Experimental Immunology, Immunology Frontier Research Center, Osaka University. 3-1 Yamadaoka, Suita, Osaka 565-0871, Japan.

## Abstract

CTLA-4 exists as membrane (mCTLA-4) and soluble (sCTLA-4) forms. Here, we show that effector-type regulatory T cells (Tregs) are main sCTLA-4 producers in basal and inflammatory states with distinct kinetics upon TCR stimulation. Mice specifically deficient in sCTLA-4 production exhibited spontaneous activation of Th1, Th17, Tfh, and Tc1 cells, autoantibody and IgE production, M1-like macrophage polarization, and impaired wound healing. In contrast, sCTLA-4-intact mCTLA-4-deficient mice, when compared with double-deficient mice, developed milder systemic inflammation and showed predominant activation/differentiation of Th2, M2-like macrophages, and eosinophils. Consistently, recombinant sCTLA-4 inhibited *in vitro* differentiation of naïve T cells towards Th1 through CD80/CD86 blockade on antigen-presenting cells, but did not affect Th2 differentiation. Moreover, sCTLA-4-intact mCTLA-4-deficient Tregs effectively suppressed Th1-mediated experimental colitis whereas double-deficient Tregs did not. Thus, sCTLA-4 production by Tregs during chronic inflammation is instrumental in controlling type 1 immunity while allowing type 2 immunity to dominate and facilitate inflammation resolution.

## Introduction

Cytotoxic T-lymphocyte-associated antigen-4 (CTLA-4) is a coinhibitory receptor that binds to CD80 and CD86 to prevent them from activating T cells via CD28. It is expressed by conventional T cells (Tconvs) only upon activation and by Foxp3-expressing CD4^+^CD25^+^ regulatory T cells (Tregs) constitutively^1^. There are two variants of CTLA-4, namely membrane and soluble CTLA-4 (mCTLA-4 and sCTLA-4, respectively), which interact with the same site of CD80/CD86. Mice with Treg-specific deficiency of both mCTLA-4 and sCTLA-4 suffer from fatal systemic autoimmunity, lymphoproliferation, and IgE hyperproduction, similar to Foxp3 mutant mice that lack Tregs^2^. In humans, CTLA-4 haplo-insufficiency evokes systemic autoimmunity/immunopathology mainly due to decreased Treg expression of CTLA-4, hence impaired Treg suppressive function^3, 4^. Single nucleotide polymorphisms in the *CTLA4* gene locus are associated with genetic risk of common autoimmune diseases such as rheumatoid arthritis, Graves’ disease, and type 1 diabetes^5, 6^. In addition, CTLA-4 blockade is effective in promoting cancer immunity, at least in part, due to reduced Treg suppressive activity, which may explain the occasional occurrences of autoimmune and immunopathological diseases as adverse effects of anti-CTLA-4 cancer immunotherapy^2, 7^. To date, mCTLA-4, rather than sCTLA-4, has been the main focus of investigation and understandably so due to its dominant role in regulating immune responses both cell-intrinsically in Tconvs and extrinsically through Tregs. However, much is still unknown about the importance of sCTLA-4, whether in physiological or disease states.

Both mCTLA-4 and sCTLA-4 express the exon2-encoded evolutionary conserved MYPPPY motif required for strong and stable binding to CD80/CD86 on APCs^8, 9^. The mCTLA-4 molecule possesses the trans-membrane domain encoded by *Ctla4* exon3 while sCTLA-4 does not. Numerous studies have shown that mCTLA-4 plays multiple roles in inhibiting activation of Tconvs, such as by competing with CD28 for binding to CD80/CD86, by transducing a negative signal into Tconvs upon interacting with CD80/CD86, or by down-regulating CD80/CD86 expressed on APCs (reviewed in ref. 1). Similarly, sCTLA-4 has been shown to modulate *in vitro* TCR-induced Tconv proliferation; and neutralization of sCTLA-4 by a specific antibody or reduction of sCTLA4 expression by siRNA knock-down promotes T-cell recall responses, decreases cancer metastasis and augments autoimmune disease development in genetically autoimmune-prone animals^10, 11^. In humans, a genetic variation that reduces sCTLA-4 transcription could predispose individuals to type 1 diabetes^5^; and the amount of sCTLA-4 in the circulation increases in various autoimmune diseases^12–15^. Despite these findings on CTLA-4, it has yet to be determined whether Tregs and Tconvs produce mCTLA-4 and sCTLA-4 in tandem upon activation and whether the two forms of CTLA-4 exert distinct immunosuppressive/immunomodulatory functions in basal and activated states.

Here, we have addressed physiological roles of sCTLA-4 in immune regulation under non-inflammatory and inflammatory conditions by generating a mouse strain specifically deficient in sCTLA-4 and another strain expressing sCTLA-4 on the mCTLA-4-deficient background. The former showed spontaneous activation of Th1 cells while the latter exhibited Th2 polarization of Tconvs that were spontaneously activated because of mCTLA-4 deficiency. Furthermore, *in vitro* CD80/CD86 blockade by sCTLA-4 inhibited naïve T cells to differentiate towards Th1, but allowed Th2 differentiation. We have also shown that activated effector Tregs are a main producer of sCTLA-4, with distinct kinetics of its production, especially in chronic inflammation and that Tregs solely producing sCTLA-4 are able to suppress experimental colitis. Thus, sCTLA-4 not only complements mCTLA-4 in immune suppression but also inhibits functional polarization of activated Tconvs to Th1 cells, allowing Th2 skewing, particularly during inflammation. sCTLA-4 can therefore be targeted to treat autoimmunity and cancer and to facilitate tissue repair.

## Results

### sCTLA-4 is predominantly produced by effector Tregs in the thymus and secondary lymphoid organs/tissues, at inflammation sites, and in tumor tissues, with distinct production kinetics upon TCR stimulation

In normal mice, both sCTLA-4 and mCTLA-4 mRNA was transcribed in the thymus solely by CD25^+^CD4^+^CD8^-^ Tregs, and in peripheral lymphoid organs by activated/effector Tregs (as CD44^hi^CD62L^lo^CD25^hi^CD4^+^ T cells) at levels ∼2-fold higher than those in CD44^lo^CD62L^hi^ resting/naïve Tregs and CD44^hi^CD62L^lo^ effector/memory CD25^-^CD4^+^ Tconvs (**Fig. 1a,b** and **Extended Data Fig. 1a-c**). There was negligible transcription of both mRNAs in CD44^lo^CD62L^hi^ naïve CD4^+^ Tconvs. Upon *in vitro* TCR stimulation, sCTLA-4 mRNA transcription was downregulated in thymic and peripheral Tregs and Tconvs (**Fig. 1c** and **Extended Data Fig. 1b,c**). In contrast, TCR stimulation steadily upregulated mCTLA-4 mRNA in both Tregs and Tconvs, resulting in a much lower ratio of sCTLA-4 mRNA to mCTLA-4 mRNA in activated Tregs and Tconvs compared to steady-state Tregs in which the amount of mRNA for sCTLA-4 was approximately 1/16 compared with mCTLA-4 mRNA (**Extended Data Fig. 1d**).

**Fig. 1:**
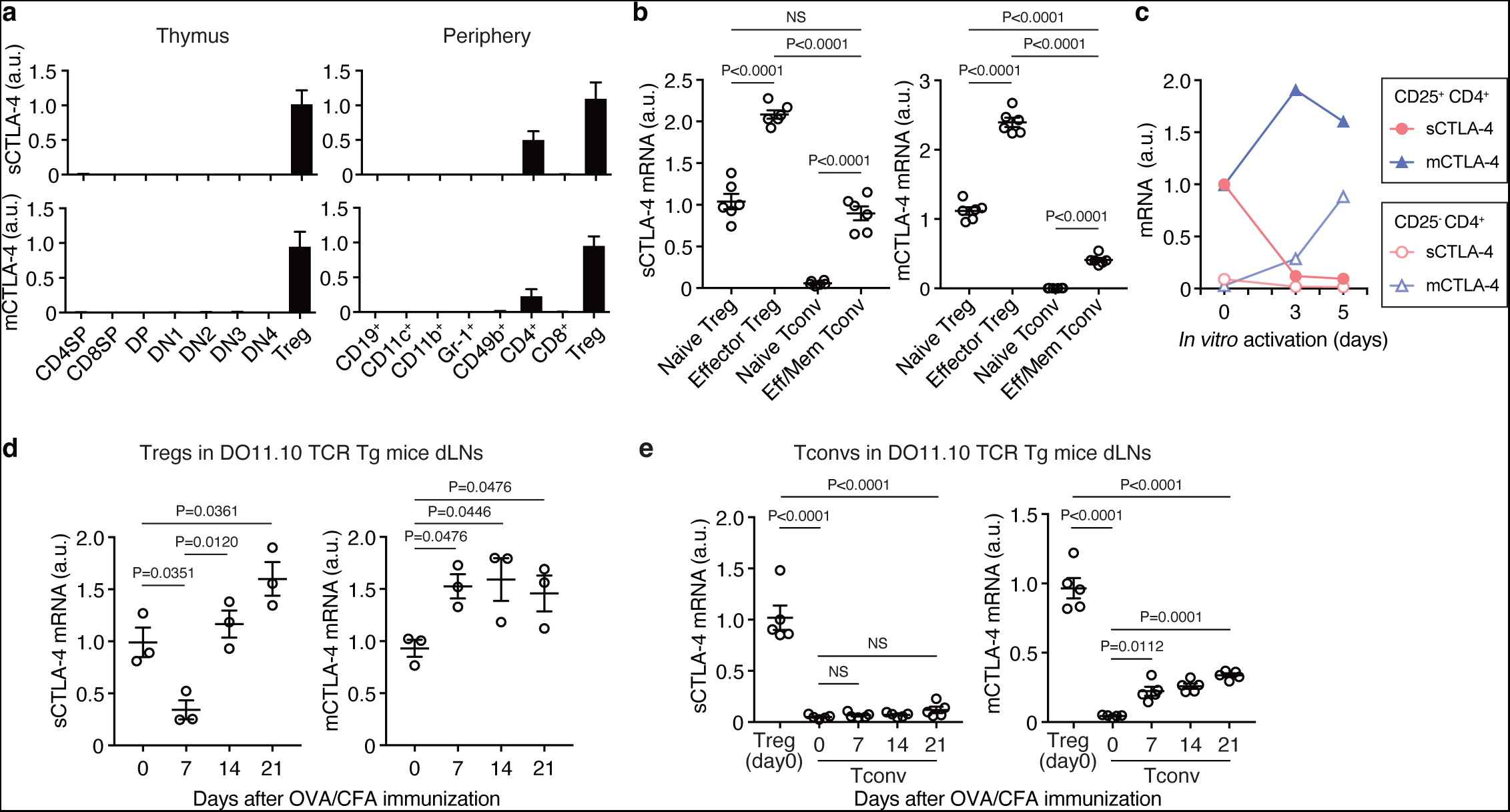
sCTLA-4 and mCTLA-4 expression by Tregs and Tconvs in steady and activated states. **a,** Predominant expression of sCTLA-4 and mCTLA-4 mRNA by Tregs in the thymus and peripheral lymphoid organs. Relative mRNA expression of sCTLA-4 and mCTLA-4 in indicated cells purified from thymi of 3-week-old BALB/c mice and peripheral lymphoid organs of 8-week-old BALB/c mice. Double negative (DN) stage of thymocytes were subdivided into 4 populations (DN1: CD44^+^CD25^-^, DN2: CD44^+^CD25^+^, DN3: CD44^-^CD25^+^, and DN4: CD44^-^CD25^-^ cells). SP, single positive; DP, double positive. **b,** Relative expression of sCTLA-4 and mCTLA-4 mRNA in indicated CD4^+^ T-cell populations sorted from 8-week- old BALB/c mice. Eff/Mem, Effector/Memory. **c,** Kinetics of relative expression of sCTLA-4 and mCTLA-4 mRNA in CD25^+^ or CD25^-^CD4^+^ T cells stimulated with anti-CD3/CD28 mAb- coated Dynabeads for indicated days. **d,e,** Relative expression of sCTLA-4 and mCTLA-4 mRNA in CD4^+^ Foxp3^+^(GFP^+^) Tregs (**d**) or CD4^+^ Foxp3^-^(GFP^-^) Tconvs (**e**) sorted from draining swollen lymph nodes (dLN) of OVA/CFA-immunized DEREG (or FDG) DO11.10 mice on indicated days after immunization. Three-week-old mice (**a**, thymocytes) or 8-week- old mice (**a**, lymphocytes, **b-e**) were used for the experiments. Error bars denote the mean ± s.e.m. in (**b,d,e**) or the mean ± s.d. in (**a,c**). Data are representative or summary of at least two independent experiments with two or more mice (**a-e**). Statistical significances were determined by one-way ANOVA with Holm-Sidak multiple comparisons test in (**b,d,e**). OVA; Ovalbumin; CFA, Complete Freund’s Adjuvant; NS, not significant.

*In vivo*, sCTLA-4 mRNA expression was downregulated in Tregs of Foxp3/GFP- reporter DO11.10 mice, which express a transgenic TCR specific for OVA peptide, until day 7 after OVA immunization. This was followed by a steady increase to above the basal level and remained high for at least 3 weeks (**Fig. 1d**). It contrasted with only a small increase in sCTLA- 4 transcription in activated Tconvs from draining lymph nodes; and the increase was to much lower levels even compared with non-stimulated Tregs (**Fig. 1e**). Both antigen-stimulated Tregs and Tconvs steadily upregulated mCTLA-4 mRNA, showing different kinetics between sCTLA-4 and mCTLA-4 production in both Tregs and Tconvs i*n vivo* as observed *in vitro* (**Fig. 1c**). Supporting the results, Tregs in male reporter mice suffering from chronic skin injuries because of fighting up-regulated the transcription of both sCTLA-4 and mCTLA-4 mRNA (**Extended Data Fig. 1e**). In addition, our analysis of public bulk RNA-seq data sets of CTLA- 4 transcription by Tregs resident in various mouse tissues (the gut, skin, and adipose tissue) revealed that tissue Tregs, a majority of which are effector-type Tregs^16^, were much more active in sCTLA-4 transcription compared with splenic or lymph node Tregs (**Extended Data Fig. 1f**). In healthy humans, CD4^+^CD45RA^-^CD25^hi^ effector Tregs in the peripheral blood showed the highest expression of sCTLA-4 mRNA among FOXP3^+^CD4^+^ T cells including CD45RA^+^CD25^+^ resting/naive Tregs and activated FOXP3^+^ non-Tregs^17^ (**Extended Data Fig. 1g**). Moreover, our analysis of a public data set of human breast cancer in which tumor-infiltrating CTLA-4^+^ Tregs and CTLA-4^+^ effector Tconvs were comparable in ratio revealed that the former transcribed sCTLA-4 mRNA much more actively than the latter (**Extended Data Fig. 2a**), as also observed in mouse tumors and intestinal tissue samples under chronic pathobiont-induced colitis (**Extended Data Fig. 2b**).

These results taken together indicate that Tregs are a main source of sCTLA-4 production in basal and activated states with transient down-regulation of the production upon TCR stimulation and subsequent up-regulation to higher levels than before.

### Specific sCTLA-4 deficiency, with intact mCTLA-4, enhances Th1, Th17, and Tfh development with autoantibody and IgE production

In order to examine sCTLA-4 functions *in vivo,* we generated a BALB/c-background mouse strain specifically deficient in sCTLA-4 with intact expression of mCTLA-4, hereafter called S^-^M^+^ mice. We constructed S^-^M^+^ BAC transgenic mice carrying a BAC sequence containing the *Ctla4* locus with conjugation of exons 3 and 4 to abrogate sCTLA-4 expression and leave the other *Ctla4* regulatory regions intact (**Fig. 2a** and **Extended Data Fig. 3a**). The mice were crossed with BALB/c S^-^M^-^ mice to solely express mCTLA-4, and a strain was selected which was equivalent in mCTLA-4 expression to WT S^+^M^+^ mice at the mRNA (**Fig. 2b**) and protein (**Extended Data Fig. 3b**) levels. Such S^-^M^+^ mice grew and survived normally as WT mice for the observation period of 30 weeks, and did not show splenomegaly or lymphadenopathy (**Fig. 2a** and **Extended Data Fig. 3c**). Histological examination, however, revealed enlarged ectopic lymphoid structures in the adipose tissue in the majority of S^-^M^+^ mice (in all the ten mice examined) and leukocyte infiltration into the lung in some (∼30%) of them (**Fig. 2c**), with no histologically evident lesions in other organs (**Extended Data Fig. 3d**). In addition, when examined at 30 weeks of age, S^-^M^+^ mice possessed significantly higher concentrations of serum IgE (**Fig. 2d**), IL-6, and TNF-α (**Fig. 2e**) than S^+^M^+^ mice. They spontaneously developed significantly high titers of serum autoantibodies, such as anti-dsDNA IgG (a marker for systemic autoimmunity) and anti-parietal cell IgG autoantibodies (a marker for organ-specific autoimmunity) (**Fig. 2f**), as seen in autoimmune diseases induced by Treg depletion^18^.

**Fig. 2:**
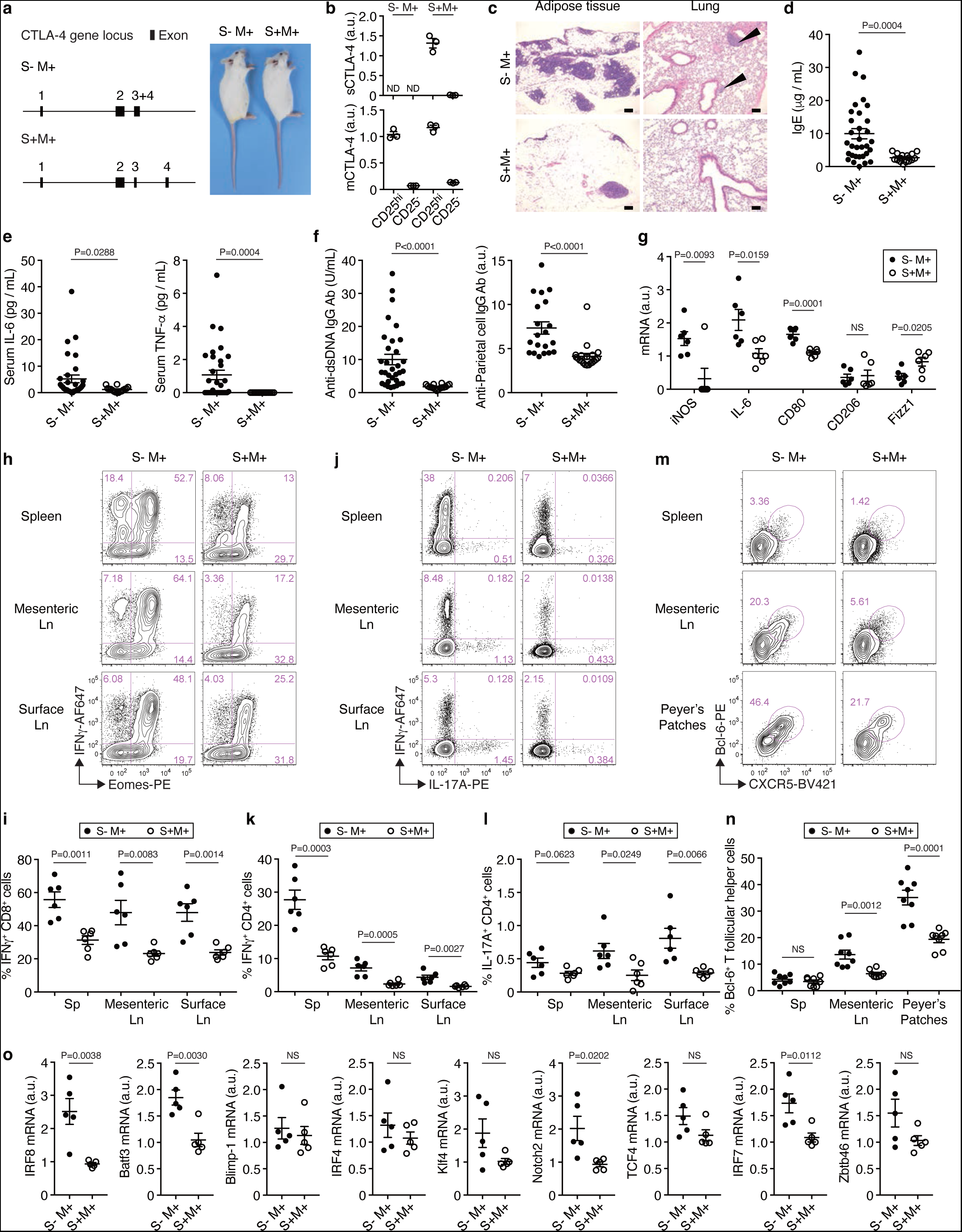
Autoimmunity with expanded/activated Th1, Tc1, Tfh, and M1-like macrophages in S^-^M^+^ mice. **a,** Schematic representation of bacterial artificial chromosome (BAC) CTLA-4 allele for preparation of S^-^M^+^ mice and a representative picture of the mice. The BAC vector RP23- 146J17 was modified to delete the intron between exon3 and exon4. The BAC transgenic mice were crossed with S^-^M^-^ mice to generate S^-^M^+^ mice. **b,** No expression of sCTLA-4 mRNA in CD4^+^ T cells from 6-week-old S^-^M^+^ mice. **c,** Histology of the adipose tissue (n = 10 mice per group) and the lung (n = 6 mice per group) in S^-^M^+^ and S^+^M^+^ mice. Scale bars represent 100 μm. **d-f,** Serum concentrations of IgE, IL-6, TNF-α, anti-dsDNA IgG antibody, and anti-parietal cell IgG antibody in indicated groups of mice. The data were accessed by Mann-Whitney test. **g,** Relative expression of mRNA for iNOS (Nos2), IL-6, CD80, CD206 (Mrc1), and Fizz1 (Retnla) in CD11b^+^ F4/80^+^ peritoneal macrophages purified from indicated groups of mice. **h-l,** Co-staining of IFNγ and Eomes in CD8^+^ T cells (**h,i**), and IFNγ and IL-17A in CD4^+^ T cells (**j-l**) of indicated groups of mice. Cells prepared from spleen, mesenteric lymph nodes, or surface lymph nodes were stimulated *in vitro* with PMA/Ionomycin/Monensin for 5 hours. **m,n,** Increase in Tfh cells as Bcl-6^+^ CXCR5^+^ Foxp3^-^ CD44^+^ CD4^+^ B220^-^ cells in mesenteric lymph nodes and Peyer’s patches from S^-^M^+^ mice. **o,** Relative expression of mRNA for major lineage transcription factors in CD11c^+^ splenic DCs purified from indicated groups of mice. Thirty-week-old mice were used for the experiments (**a,c-o**). All error bars denote the mean ± s.e.m. Data are representative or summary of at least two independent experiments with three or more mice per group (**b-o**). Statistical significances were determined by unpaired two-tailed *t*-test in (**g,i,k,l,n,o**) or Mann-Whitney test (**d-f**). BV421, Brilliant Violet 421; AF647, Alexa Fluor 647; dsDNA, double stranded DNA; ND, not detected.

The proportion of CD44^+^CD8^+^ T cells gradually increased with age in the spleen and lymph nodes, for example, with more than 2-fold increase at 24 weeks compared with at 8 weeks of age (**Extended Data Fig. 3e**), accompanied by a smaller but significant increase in CD44^+^CD4^+^ T cells in lymph nodes (**Extended Data Fig. 3f**). Moreover, 30-week-old S^-^M^+^ mice showed a marked expansion of IFNγ^+^ cells among CD8^+^Eomes^+^ effector/memory T cells and among CD4^+^ T cells in the spleen, mesenteric and other lymph nodes (**Fig. 2h-k**). IL- 17A^+^CD4^+^ T cells also increased significantly in S^-^M^+^ mice (**Fig. 2j,l**).

S^-^M^+^ mice harbored an increased proportion of Bcl-6^+^PD-1^+^CXCR5^+^ Tfh cells in Peyer’s patches and mesenteric lymph nodes (**Fig. 2m,n** and **Extended Data Fig. 3g).** The number of IL-4-secreting Tfh cells significantly increased (∼five-fold) in Peyer’s patches, although the ratio among Tfh cells was not significantly different (**Extended Data Fig. 3h,i**). The mice also possessed an increased proportion of CD38^-^GL7^+^IgD^-^IgM^-^ germinal center B cells in mesenteric lymph nodes (**Extended Data Fig. 3j**). In addition, the high proportion of Tfh and Th1 cells in the Peyer’s patches and mesenteric lymph nodes was accompanied by a relative decrease in the proportion of Tregs (**Extended Data Fig. 3k-m**). Thus, with scarcely detectable GATA-3^+^ Th2 cells in lymph nodes (data not shown), the hyper-IgE in S^-^M^+^ mice could be attributed to the expansion of IL-4^+^ Tfh cells, which induced IgE class switching in B cells^19^, in the gut-associated lymphoid tissue.

qPCR analysis of peritoneal cavity macrophages isolated from S^-^M^+^ mice showed higher expression of M1 macrophage markers such as iNOS, IL-6, and CD80, with lower expression of M2 macrophage markers such as CD206 (Mannose Receptor C-type 1) and Fizz1, indicating M1-like macrophage polarization in S^-^M^+^ mice (**Fig. 2g**). DCs in aged S^-^M^+^ mice were higher than those in S^+^M^+^ mice in the expression of IRF8, Batf3, and Notch2 (**Fig. 2o**), suggesting a predominance of cDC1/Notch2^+^cDCs and their possible contributions to Th1 and Th17 induction^20^.

Taken together, the results indicate that sCTLA-4 physiologically suppresses T-cell differentiation into Th1, Th17, Tfh, and Tc1 cells, and also prevents M1-like macrophage polarization and cDC1 differentiation, and that specific sCTLA-4 deficiency spontaneously evokes marked type 1 immune responses that can result in mild autoimmunity with aging in otherwise normal mice.

### sCTLA-4 ameliorates and modulates systemic inflammation in sCTLA-4-intact mCTLA- 4-deficient mice

Next, to further assess sCTLA-4 function *in vivo*, we generated a BALB/c-background knock-in mouse stain that expressed sCTLA-4 but not mCTLA-4 (abbreviated as S^+^M^-^ mice) by deleting the splicing acceptor site upstream of *Ctla4* exon3 and leaving the other *Ctla4* regulatory regions the same as in WT mice (**Fig. 3a** and **Extended Data Fig. 4a**). Peripheral CD4^+^ T cells in S^+^M^-^ mice indeed expressed sCTLA-4 without mCTLA-4 at the protein and mRNA levels in the steady state and after *in vitro* activation (**Fig. 3b** and **Extended Data Fig. 4b**). The sCTLA-4 concentration in S^+^M^-^ mice (average 1.5 ng/ml), detectable by ELISA (with CTLA-4 pan-specific mAb), was significantly higher than in WT mice (**Fig. 3c**), being equivalent or lower compared to human autoimmune disease patients (e.g., 10-30 ng/ml in SLE^15^ or Graves’ disease^13^ with <5 ng/ml in healthy individuals). Thus, the high sCTLA-4 concentration in S^+^M^-^ mice likely reflected Treg and Tconv activation and their numerical increases in tissues and the blood due to systemic autoimmunity (see below), which could upregulate CTLA-4 transcription/expression (**Fig. 1d**). In addition, the loss of the exon3 splicing acceptor site in S^+^M^-^ mice appeared to make mRNA for mCTLA-4 to be transcribed as sCTLA-4 mRNA in both activated Tregs and Tconvs, contributing to the high serum sCTLA- 4 concentration in these mice.

**Fig. 3:**
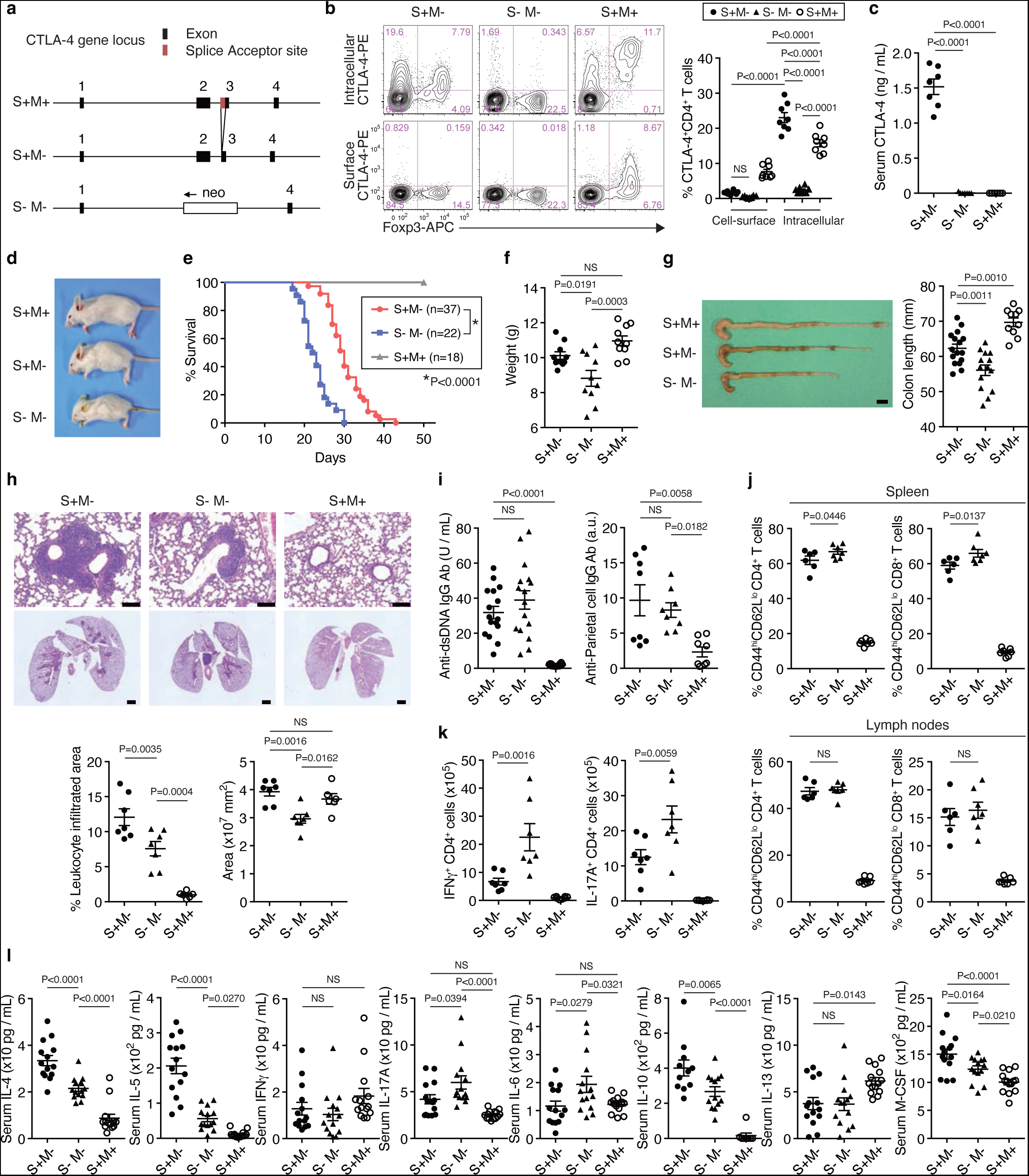
S^+^M^-^ mice are immunologically distinct from S^-^M^-^ mice. **a,** Schematic representations of the CTLA-4 gene locus of S^+^M^+^, S^+^M^-^, and S^-^M^-^ mice. Splice acceptor site (86 bp) in front of exon3 was deleted in S^+^M^-^ mice. **b,** Co-staining of CTLA-4 and Foxp3 in CD4^+^ T cells from indicated groups of 3-week-old mice after stimulation of PMA/Ionomycin/Monensin for 4 hours. **c**, Serum concentrations of sCTLA-4 in indicated groups of mice. **d,** Systemic appearance of the mice in (**a**). **e,** Survival of S^+^M^-^, S^-^M^-^, and S^+^M^+^ mice. Statistical significance between S^+^M^-^ and S^-^M^-^ mice was accessed by Log-rank test. **f,** Body weight of the mice in (**a**). **g,** Colon lengths of indicated groups of mice. Scale bar represents 5 mm. **h**, Lung inflammation in S^+^M^-^ and S^-^M^-^ mice. Hematoxylin and eosin (HE) staining of the lung tissues of indicated groups of mice. Percentages of leukocyte infiltrated areas (leukocyte foci excluding lymph nodes > 1,000 μm2) among whole areas of lung sections without cavities were determined by computational analysis. Scale bars represent 100 μm in upper figures and 1,000 μm in lower figures. **i,** Serum titers of anti-dsDNA IgG antibody and anti-parietal cell IgG antibody in indicated groups of mice. **j,** Increase in CD44^hi^CD62L^lo^ CD4^+^ and CD8^+^ T cells in S^+^M^-^ and S^-^M^-^ mice. **k,** Suppression of IFNγ- or IL-17A-producing CD4^+^ T-cell expansion in mesenteric lymph nodes of S^+^M^-^ mice. **l,** Serum concentrations of cytokines in indicated groups of mice. Three-week-old mice were used for the experiments in (**a-l**) because of the early mortality of S^-^M^-^ mice. Error bars denote the mean ± s.e.m. in (**b,c,f-l**). Data are representative or summary of at least three independent experiments with six or more mice (**b-l**). Statistical significances were determined by one-way ANOVA with Holm-Sidak multiple comparisons test in (**b,c,f-l**).

Compared with S^-^M^-^ mice, which developed fatal systemic autoimmunity^21, 22^, S^+^M^-^ mice were less debilitated (**Fig. 3d**) and had significantly longer lifespan (**Fig. 3e**). Compared with S^-^M^-^ mice at 3 weeks of age, S^+^M^-^ mice exhibited less body weight loss (**Fig. 3f**) and less severe spontaneous colitis as evidenced by milder shrinkage of the colon (**Fig. 3g**) that contained less inflammatory cell infiltrates (**Extended Data Fig. 4c**). However, we found that the severity of inflammation varied from organ to organ; for example, the lungs of S^+^M^-^ mice were more inflamed with larger areas of leukocyte infiltration than the lungs of S^-^M^-^ mice (**Fig. 3h**). Both strains exhibited cellular infiltration into the pancreas, liver, salivary gland, heart, and gastric mucosa, with various extents of tissue damage (**Extended Data Fig. 4d**). In correlation with the tissue damage, they showed elevated serum amylase, AST, ALT, and bilirubin, and decreased serum albumin (**Extended Data Fig. 4e**). They also suffered from anemia (**Extended Data Fig. 4f**). Serum titers of anti-dsDNA and anti-gastric parietal cell autoantibodies were comparable between S^+^M^-^ and S^-^M^-^ mice (**Fig. 3i**). No significant differences were found in serum concentration of each class of immunoglobulin (**Extended Data Fig. 4g**).

S^+^M^-^ and S^-^M^-^ mice similarly developed splenomegaly and lymphadenopathy with increases in CD44^+^CD62L^low^ activated CD4^+^ and CD8^+^ T cells (**Fig. 3j** and **Extended Data Fig. 4h**). The lymphocyte cellularity of lymph nodes was much lower in the former than the latter, with equal increases in the spleen (**Extended Data Fig. 4i-k)**. Both strains showed significant increases in the number of Tregs in lymph nodes, with their slight increases in the spleen (**Extended Data Fig. 4l**). Of note, however, is that the number of Th1 and Th17 cells was significantly lower in the mesenteric lymph nodes of S^+^M^-^ mice (**Fig. 3k**). Furthermore, serum concentrations of cytokines significantly varied between S^+^M^-^ and S^-^M^-^ mice; for example, the former were significantly higher than the latter in IL-4, IL-5, IL-10, and M-CSF, while significantly lower in IL-6 and IL-17A, with no significant differences in IFNγ and IL- 13 (**Fig. 3l**).

Collectively, sCTLA-4 was able to partially alleviate the severe systemic autoimmunity in S^-^M^-^ mice, although it was unable to fully compensate for the loss of mCTLA-4 in S^+^M^-^ mice, which underscored a critical role of mCTLA-4 in immunological self-tolerance. sCTLA- 4 might also play a specialized role in immune regulation as manifested by the difference between S^+^M^-^ and S^-^M^-^ mice in the cytokine production patterns, skewing in Th cell composition, and the inflammation phenotype in some organs (e.g., the lung).

### sCTLA-4 promotes M2-like macrophage polarization during chronic inflammation

The above results with S^-^M^+^ and S^+^M^-^ mice suggested that sCTLA-4 might be immunomodulatory as well as immunosuppressive through its effect on the composition and/or function of CD80/CD86-expressing immune cells, such as CD11b^+^ cells (mainly monocytes/macrophages and granulocytes), CD11c^+^ cells [dendritic cells (DCs)], and B220^+^ cells (B cells).

We first noted that, despite severe systemic inflammation in both S^-^M^-^ and S^+^M^-^ strains, the latter did not show a significant increase (compared with S^+^M^+^ mice) in the proportion of CD80^+^CD11b^+^ cells among splenic CD11b^+^ cells, contrasting with a marked increase in S^-^M^-^ mice (**Fig. 4a** and **Extended Data Fig. 5a**). The results suggested that sCTLA-4 binding to CD80 *in vivo* might hinder *ex vivo* anti-CD80 mAb staining of CD80^+^CD11b^+^ cells, hence their apparent reduction, in S^+^M^-^ mice. The possibility, however, could be excluded by the following findings. First, the frequencies of CD80^+^ and CD86^+^ cells among CD11c^+^ and B220^+^ cells did not differ significantly between the two strains. Second, the CD80 mRNA level in CD11b^+^ cells from S^+^M^-^ mice was significantly lower compared with those from S^-^M^-^ mice, whereas the CD86 mRNA level was not (**Extended Data Fig. 5b**). Further, anti-CD80 mAb (16-10A1) used for cell staining blocked binding of CTLA4-Ig to CD80^23^, albeit the binding was of higher avidity than that of sCTLA-4 (see below).

**Fig. 4:**
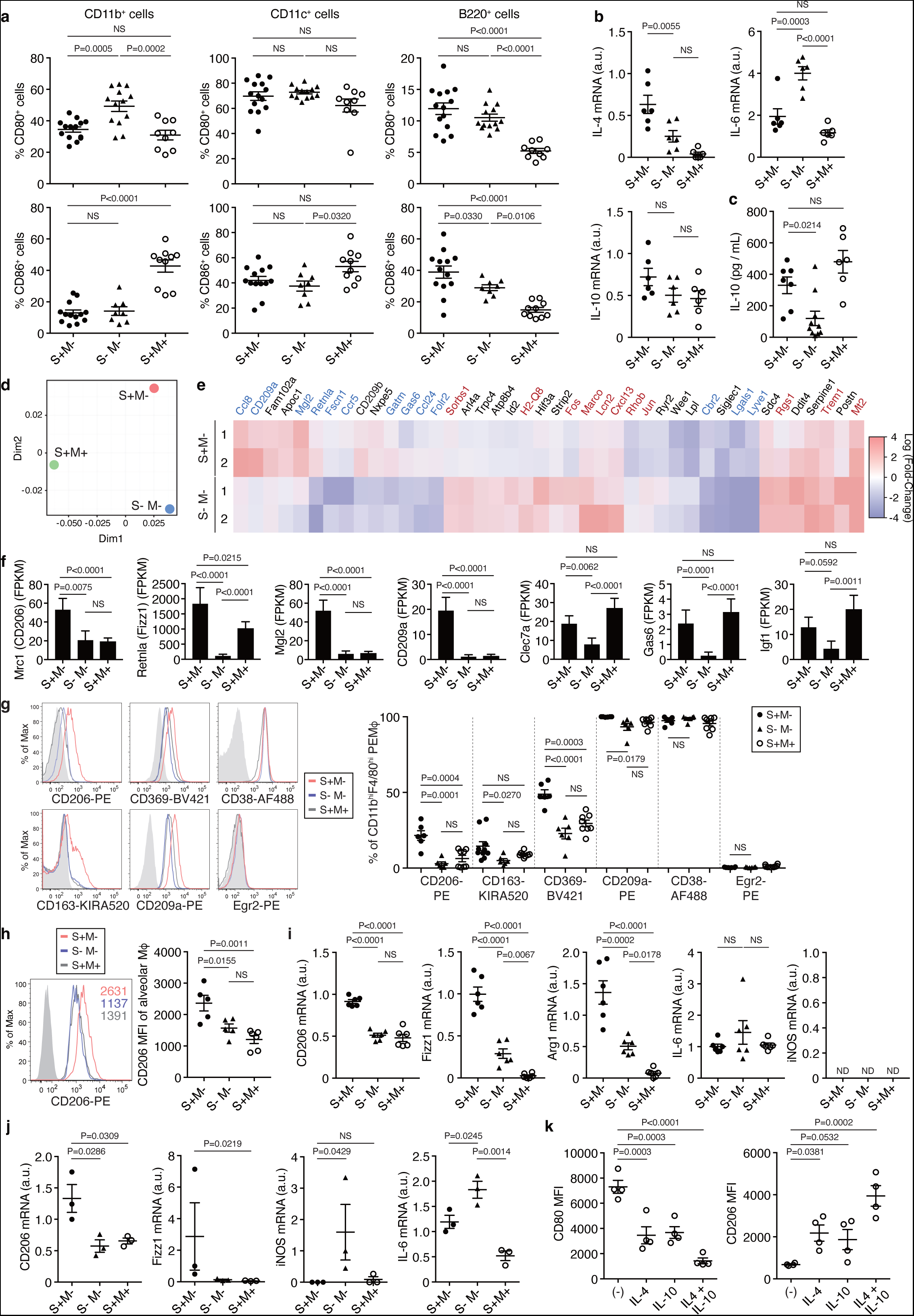
M2-like macrophage polarization in S^+^M^-^ mice. **a,** CD80 and CD86 expression by splenic CD11b^+^, CD11c^+^, and B220^+^ cells in indicated groups of mice. **b,** Relative expressions of IL-4, IL-6, and IL-10 mRNA in splenic CD11b^+^ cells purified from indicated groups of mice. **c,** Concentration of IL-10 in culture supernatants taken 24 hours after *in vitro* culture of F4/80^+^CD11b^+^ peritoneal macrophages from indicated groups of mice. **d,** Multi-dimensional scaling plot (PCoA) of RNA-seq transcriptomes of F4/80^+^CD11b^+^ peritoneal macrophages from indicated groups of mice (n = 2 per group). **e,** Heatmap displaying fold change in gene expression by peritoneal macrophages from S^+^M^-^ and S^-^M^-^ mice compared with S^+^M^+^ mice (genes with FDR-adjusted *P* < 0.001: S^+^M^-^ versus S^-^M^-^). Red, M1 macrophage markers. Blue, M2 macrophage markers. **f,** Expression of typical genes induced by IL-4/IL-13 in macrophages. The levels of mRNA were determined by RNA-seq analysis. Statistical significances denote FDR-adjusted *P* values for multiple testing. **g,** Significant increase in CD206 (Mrc1), CD163, CD369 (Dectin-1, Clec7a), and CD209a (DC- SIGN) expressing peritoneal macrophages in S^+^M^-^ mice determined by flow cytometry. **h,** Increase in CD206 expression on alveolar macrophages defined as CD11c^+^Siglec- F^+^CD11b^int^CD64^+^ cells in S^+^M^-^ mice. MFI, mean fluorescence intensity. **i,** Relative expression of mRNA for CD206, Fizz1 (Retnla), Arg1, IL-6, and iNOS (Nos2) in alveolar macrophages purified from indicated groups of mice. **j,** Relative expression of mRNA for CD206, Fizz1, iNOS, and IL-6 in colonic CD45^+^Siglec-F^-^Ly6G^-^CD11b^+^F4/80^+^ macrophages purified from indicated groups of mice. The data of Fizz1 and iNOS were accessed by Kruskal-Wallis test followed by Dunn’s test. **k,** CD80 and CD206 MFIs of splenic CD45^+^CD68^+^F4/80^+^ cells in S^-^ M^-^ mice 24 hours after treatment with IL-4 and IL-10 on a culture dish immobilized with hydrophilic polymer. Three-week-old mice were used for all the experiments (**a-k**). Error bars denote the mean ± s.e.m. in (**a-c,g-k**). Data are representative or summary of at least two independent experiments with three or more mice per group (**a-c,g-k**), or RNA-seq experiments with two mice per group (**d-f**). Statistical significances were determined by one-way ANOVA with Holm-Sidak multiple comparisons test in (**a-c,g-k**). FDR, False Discovery Rate; AF488, Alexa Fluor 488; KIRA520, KIRAVIA Blue 520.

Next, to determine whether sCTLA-4 altered the phenotype and function of CD11b^+^ cells, we analyzed their cytokine expression patterns in the spleen and peritoneal cavity by qPCR. Splenic CD11b^+^ cells in S^+^M^-^ mice were significantly higher than those in S^-^M^-^ or S^+^M^+^ mice in the transcription of IL-4, with no significant difference in IL-10 transcription, while CD11b^+^ cells in S^-^M^-^ mice were higher in IL-6 transcription compared with other strains (**Fig. 4b**). CD11b^+^F4/80^+^ peritoneal macrophages, a dominant population of CD11b^+^ cells in the peritoneal cavity, in S^+^M^-^ mice showed higher *in vitro* production of IL-10 than those in S^-^M^-^ mice, with comparable numbers of the macrophages in these strains (**Fig. 4c**).

RNA-seq of CD11b^+^ peritoneal macrophages purified from S^+^M^-^, S^-^M^-^, or S^+^M^+^ mice further revealed that the macrophages in S^+^M^-^ mice were distinct from those in S^-^M^-^ mice (**Fig. 4d**), especially in the expression of the genes characterizing M1- or M2-like macrophages in terms of their functions of phagocytosis, wound healing, cell chemotaxis, and angiogenesis (**Fig. 4e** and **Extended Data Fig. 5c**). For example, peritoneal macrophages in S^+^M^-^ mice highly expressed M2 markers such as *Mrc1* (CD206), *Retnla* (Fizz1), *Mgl2*, *CD209a* (encoding DC-SIGN), *Clec7a* [encoding Dectin-1 (CD369)], *Gas6, Igf1*, *Arg1*, and *Ym1*, whereas S^-^M^-^ mice macrophages highly expressed M1 markers such as *NOS2* (iNOS), *Il6*, and *Tnf* (**Fig. 4f** and **Extended Data Fig. 5d,e**). KEGG pathway analysis also indicated that Th17 pathway-related genes (e.g., IL-6, TNF-α, and COX2) were profoundly down-regulated, while the genes (e.g., MMP9) related to tissue remodeling were up-regulated, in S^+^M^-^ macrophages compared with S^-^M^-^ macrophages (**Extended Data Fig. 5d**). Flow cytometric analysis of S^+^M^-^ peritoneal macrophages indeed showed significantly higher expressions of M2 markers such as CD369, CD209a, and also CD206 and CD163, both of which are common M2 markers that are relatively independent of tissue localization and macrophage origins (**Fig. 4g**). There was no difference in the expression of CD38 (M1 marker) and Egr2 (M2 marker), both of which were upregulated in bone marrow-derived macrophages *in vitro*^24^. Although alveolar macrophages have a basic M2-like phenotype, S^+^M^-^ alveolar macrophages showed a more M2-like polarization with significantly increased expressions of CD206, Fizz1, and Arg1 (**Fig. 4h,i**).

In gut-associated lymphoid tissues, where sCTLA-4 was significantly more abundant than in other lymph nodes in normal mice (**Extended Data Fig. 5f**), colonic macrophages isolated from S^+^M^-^ mice were higher than those from S^-^M^-^ mice in the expression of CD206 and Fizz1, while the latter were much higher in the expression of iNOS and IL-6 (**Fig. 4j**). S^+^M^-^ mice also harbored a larger proportion of Ly6C^lo^ circulating monocytes, which tend to differentiate into M2-like macrophages, and a lower proportion of Ly6C^hi^ monocytes, which are prone to differentiate into M1-like macrophages^25, 26^ (**Extended Data Fig. 5g**), in accord with high serum concentration of M-CSF, a cytokine required for maintaining Ly6C^lo^ monocytes^27^ (**Fig. 3l**).

Macrophages isolated from either S^+^M^-^ or S^-^M^-^ mice had similar expression levels of *Il4ra*, *Il13ra1*, *Il10ra*, and *Il10rb* receptors (**Extended Data Fig. 5e**), suggesting that the phenotypic differences in macrophages between S^+^M^-^ and S^-^M^-^ mice could be attributed to the abundance of Th2 cytokines, not to cytokine receptors expression, in the former. *In vitro* IL-4 or IL-10 treatment, particularly in combination, indeed decreased CD80 expression and increased CD206 expression in splenic macrophages isolated from S^-^M^-^ mice (**Fig. 4k**), mimicking the CD80^lo^ and CD206^hi^ phenotype of macrophages in S^+^M^-^ mice. In addition, IL- 4 and/or IL-10 (but not IL-5) induced the M2-like macrophage phenotype (e.g., CD206 and Arg1 expression) and suppressed M1 marker (e.g., iNOS) expression in RAW264.7, a macrophage cell-line. This was, however, not the case for sCTLA-4, indicating that sCTLA-4 was unable by itself to induce M2-like macrophages (**Extended Data Fig. 5h,i**).

Taken together, under a chronic autoimmune condition in S^+^M^-^ mice, secreted sCTLA-4 could alter the production of IL-4, IL-10, and IL-6 by CD11b^+^ monocytes/macrophages/granulocytes. It is plausible that the IL-4 and IL-10 produced from CD11b^+^ cells as well as those from other cell sources (see below), drove macrophages to differentiate into M2-like macrophages with the CD206^hi^CD80^lo^ phenotype.

### sCTLA-4 facilitates Th2 differentiation and eosinophil expansion under mCTLA-4- deficient chronic inflammation

We next analyzed the effects of sCTLA-4 on the cytokine profiles of other immune cells in the circulation of S^+^M^-^ mice. In accord with their high serum concentration of IL-4 and IL-5, but not IFNγ (**Fig. 3l**), S^+^M^-^ mice harbored significantly higher frequencies of IL-4 and/or IL-5-producing circulating CD4^+^ T cells than S^-^M^-^ mice (**Fig. 5a**), with no significant differences in the frequency of IFNγ-producing cells (data not shown).

**Fig. 5:**
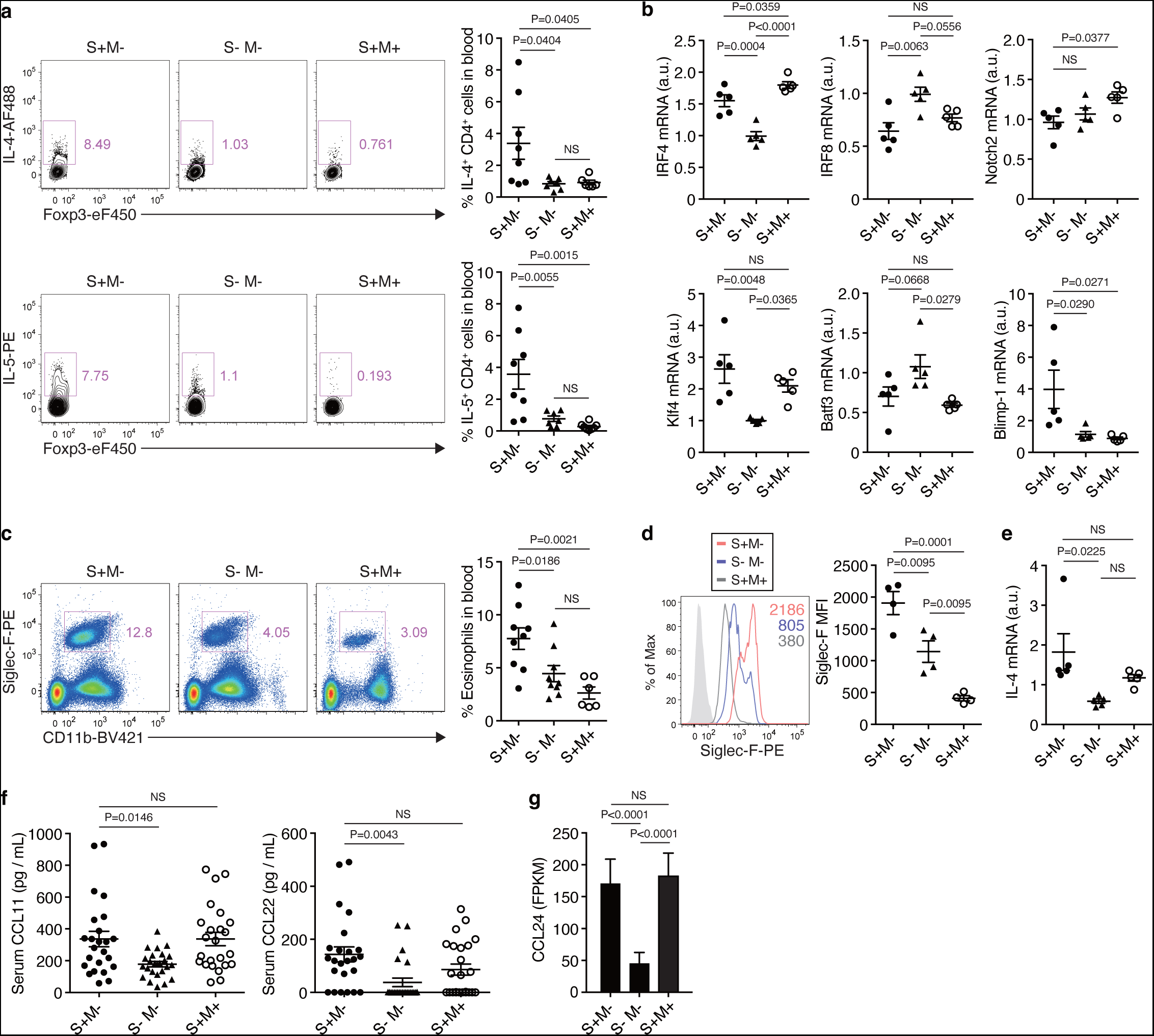
Expansion of Th2 and eosinophils, and production of Th2-type chemokines in S^+^M^-^ mice. **a,** Frequencies of IL-4- and IL-5-producing cells in Foxp3^-^CD4^+^ blood cells prepared from indicated groups of mice and *in vitro* stimulated with PMA/Ionomycin/Monensin for 5 hours. **b,** Relative expression of mRNA for major lineage transcription factors in CD11c^+^ splenic DCs purified from indicated groups of mice. **c,** Frequency of eosinophils as CD11b^int^ Siglec-F^+^ cells among CD45^+^ blood cells in indicated groups of mice. **d,** Siglec-F expression of splenic eosinophils in indicated groups of mice. MFI, median fluorescence intensity. **e,** Relative expression of IL-4 mRNA in splenic eosinophils in indicated groups of mice**. f,** Serum concentrations of CCL11 (Eotaxin-1) and CCL22 in each group of mice. **g,** CCL24 (Eotaxin- 2) mRNA expression by peritoneal macrophages in indicated groups of mice. Statistical significances denote FDR-adjusted *P* values for multiple testing. Three-week-old mice were used for all the experiments in (**a-g**). Error bars denote the mean ± s.e.m. in (**a-f**). Data are representative or summary of at least two independent experiments with four or more mice per group (**a-f**), or RNA-seq experiments with two mice per group (**g**). Statistical significances were determined by one-way ANOVA with Holm-Sidak multiple comparisons test in (**a-f**). eF450, eFluor 450.

Despite no significant differences between S^+^M^-^ and S^-^M^-^ mice in CD11c^+^ DCs in terms of the intensity of CD80/CD86 expression and the proportion of CD80/CD86-expressing cells (**Fig. 4a**), gene expression profiling revealed that DCs in S^+^M^-^ mice were higher in the expression of IRF4, Klf4, and Blimp1 and lower IRF8 and Batf3 expression, while Notch2 expression was comparable between the two strains (**Fig. 4b**), indicating a predominance of cDC2 over cDC1 in S^+^M^-^ mice^20^.

The proportion of eosinophils, as measured as CD11b^+^Siglec-F^+^ cells, was significantly higher in the blood of S^+^M^-^ mice (**Fig. 5c**). Mature eosinophils infiltrating into the spleen in S^+^M^-^ mice upregulated Siglec-F (**Fig. 5d**), and expressed IL-4 mRNA more than 2- fold as compared to eosinophils in S^-^M^-^ mice (**Fig. 5e**). Th2-associated eotaxins (i.e., CCL11 and CCL24) and CCL22^28–32^ were significantly higher in S^+^M^-^ mice than in S^-^M^-^ mice in terms of serum concentration (**Fig. 5f**) and gene transcription in peritoneal macrophages (**Fig. 5g**).

Since type 2 innate lymphoid cells (ILC2), basophils, and mast cells are also able to produce IL-4 and/or IL-5, we assessed in S^+^M^-^ mice the proportions of these cell populations in the circulation and measured the serum concentrations of IL33 and TSLP, which promote them to proliferate and produce IL-4 or IL-5^33–38^. In comparison to S^+^M^+^ mice, the proportions of whole Lin^-^CD45^+^CD90^+^ ILCs in the blood and the mesentery of S^+^M^-^ and S^-^M^-^ mice were significantly decreased (**Extended Data Fig. 6a,b**), with a non-detectable number of GATA- 3^+^ ILC cells (data not shown). Basophils, identified as CD200R3^+^c-kit^-^ cells, and c-kit^+^ mast cells were unchanged in S^+^M^-^ mice (**Extended Data Fig. 6c**). Serum IL-33 measurement showed a decreasing trend, in both S^+^M^-^ and S^-^M^-^ mice compared with S^+^M^+^ mice, while these groups of mice showed no significant difference in the serum TSLP level (**Extended Data Fig. 6d**).

These results collectively indicate that the abundance of sCTLA-4 produced in S^+^M^-^ mice under chronic inflammation blocks CD80/CD86 on APCs including DCs, thereby contributing to T-cell differentiation into IL-4/IL-5-producing Th2 cells. Th2 cells in turn augmented eosinophil migration and IL-4 production in accord with the roles of Th2 cytokines, especially IL-5, in driving activation and maturation of eosinophils to express more IL-4^39, 40^ and their signature gene Siglec-F^41^. Furthermore, IL-4 from Th2 cells, macrophages, and eosinophils, together with IL-10 from Th2 cells and macrophages, polarize the phenotype and function of macrophages to M2-like in S^+^M^-^ mice.

### sCTLA-4 inhibits Th1 but allows Th2 polarization *in vitro*

In order to determine the manner by which sCTLA-4 controlled Th1/Th2 differentiation, we assessed the effects of sCTLA-4-dependent CD80/CD86 blockade on *in vitro* activation and differentiation of WT naïve Tconvs into Th1 and Th2 cells. We prepared recombinant WT sCTLA-4 and a mutant form with Y139A alteration, which hindered its binding to CD80/CD86. The recombinant WT sCTLA-4 specifically recognized CD80/CD86^+^ CD11c/CD11b^+^ cells and pulled down CD80/86 molecules by immunoprecipitation, which was abrogated by CTLA4-Ig (CTLA-4/IgG2a Fc chimera) co-incubation (**Extended Data Fig. 7a-d**). WT sCTLA-4 was thus weaker in its avidity for CD80/86 than CTLA4-Ig because the former was monomer^42^, the latter dimer. The results were consistent with the data by surface plasmon resonance analysis, which revealed that monomeric CTLA-4 bound only with low affinity and rapidly dissociated from CD80/CD86 while dimeric CTLA-4 showed high avidity binding with decreased dissociation^43^.

*In vitro* anti-CD3 mAb stimulation of naive CD4^+^ T cells in the presence of T cell-depleted spleen cells as APCs drove CD4^+^ T cells to differentiate into IFNγ-secreting cells without exogenous addition of cytokines because of spontaneous production of IL-12 by APCs, even with Th2-prone BALB/c T cells and APCs^44, 45^. WT sCTLA-4 inhibited the proliferation of naïve CD4^+^ T cells more potently than Y139 mutant but less potently than CTLA4-Ig at the same concentration and number of APCs (**Fig. 6a**). Furthermore, WT sCTLA-4 and CTLA4- Ig profoundly hampered spontaneous Th1 differentiation in a dose-dependent fashion, while Y139A sCTLA-4 did not (**Fig. 6b,c,e**). Under a Th2-inducing condition with IL-4, recombinant WT sCTLA-4 allowed Th2 differentiation with a dose-dependent increase in Th2 cells (**Fig. 6b,d,e**).

**Fig. 6:**
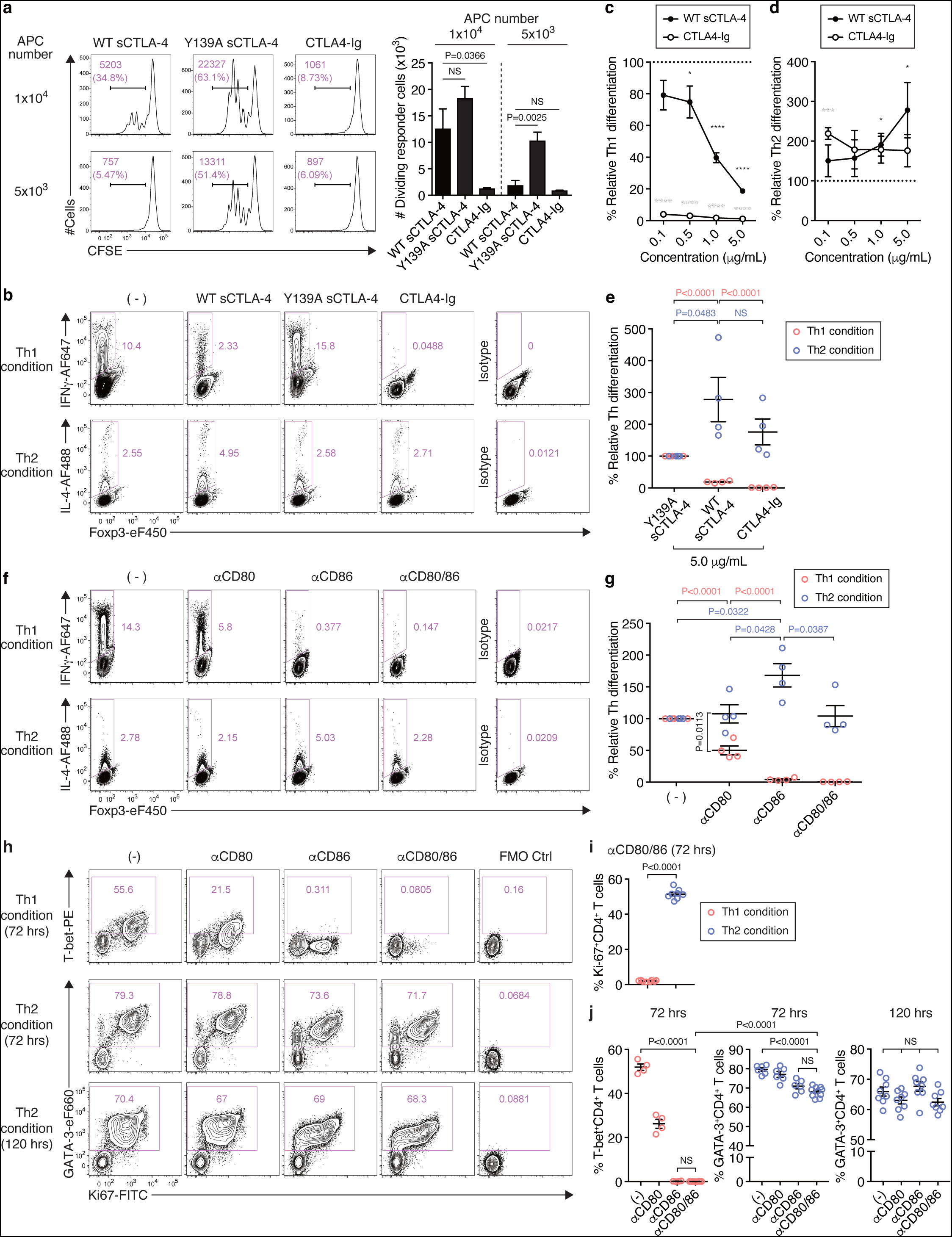
sCTLA-4 inhibits Th1 but allows Th2 differentiation *in vitro*. **a**, Effects of sCTLA-4 on *in vitro* proliferation of Tconvs. 5x104 CD45.1^+^CD4^+^CD25^-^CD62L^+^ cells were stimulated with anti-CD3 mAb in the presence of CD45.2^+^ splenocytes as APCs and 1 μg/mL sCTLA-4 recombinant protein. One-way ANOVA with Holm-Sidak multiple comparisons test. **b,** Th1 and Th2 differentiation in the presence of sCTLA-4 recombinant protein. 5x104 CD4^+^CD25^-^CD62L^+^ cells were stimulated with anti-CD3 mAb and 1x105 T-cell depleted splenocytes for 3 days with 5.0 μg/mL WT or Y139A mutant sCTLA-4, or CTLA4- Ig and then stimulated with PMA/Ionomycin/Monensin for 5 hours. **c,d,** Dose-dependent effects of WT sCTLA-4 and CTLA4-Ig on the differentiation of naive T cells towards Th1 (**c**) and Th2 (**d**). The ratio of Th1 or Th2 differentiation was calculated based on the effects of Y139A sCTLA-4 as negative control (100% horizontal dotted line). \**P* < 0.05, \*\*\**P* < 0.001, \*\*\*\**P* < 0.0001, compared with Y139A sCTLA-4 (unpaired two-tailed *t*-test). **e,** Ratios of skewed Th1 or Th2 cells in the presence of indicated recombinant proteins at 5.0 μg/mL as described in (**c**) and (**d**). One-way ANOVA with Holm-Sidak multiple comparisons test. **f,** Th1 and Th2 differentiation in the presence of CD80/86 blocking antibodies, as shown in (**b**). **g,** Ratios of skewed Th1 or Th2 cells as shown in (**f**). Ratios were calculated based on the condition without CD80/CD86 blockade. One-way ANOVA with Holm-Sidak multiple comparisons test. **h-j,** Effect of αCD80/86 antibody on T-bet^+^ Th1 differentiation and GATA-3^+^ Th2 differentiation. αCD80/86 antibody inhibits Th1 proliferation (**h,i**) and differentiation (**h,j**) compared to Th2. Two-way ANOVA (with Holm-Sidak multiple comparisons test) to analyze the effects of factor “CD80/86 availability” and factor “condition/differentiation (Th1 or Th2)” revealed statistical significances in the former factor (*P* < 0.0001), the latter factor (*P* < 0.0001), and the interaction (*P* < 0.0001). All error bars denote the mean ± s.e.m. Data are representative or summary of three independent experiments (**a**) or four or more independent experiments (**b-j**). Data points show the results of experiments with cells isolated from different mice (n = 4 to 10 mice per experiment). eF660, eFluor 660.

To confirm the above findings with sCTLA-4, we added blocking anti-CD80 or anti- CD86 mAb, or both, to anti-CD3-stimulated co-culture of naive CD4^+^ T cells and APCs with or without IL-4 (**Fig. 6f,g**). This blockade of CD80 or CD86, especially the latter, significantly hindered Th1 differentiation in the absence of IL-4, but allowed Th2 differentiation in the presence of IL-4. Cell staining for Ki67 together with T-bet or GATA-3 revealed that the blockade, especially of CD86, strongly inhibited the development of Ki67^+^T-bet^+^ cells but not of Ki67^+^GATA-3^+^ cells; the latter differentiation was not affected by prolonged (120 hours) blockade (**Fig. 6h-j**). Cell staining for Nur77 and CD69 to assess TCR signal intensity and cell proliferation showed that Nur77 expression was similarly induced under both Th1 and Th2 conditions and significantly suppressed by CD86 or CD80/CD86 blockade for 24 hours (**Extended Data Figure 7e,f**). Notably, the blockade more strongly inhibited cell proliferation of Ki67^+^Nur77^+^ cells in the Th1 condition compared with the Th2 condition, despite more suppressed Nur77 expression in the Th2 condition (**Extended Data Figure 7g,h**). Even under CD80/CD86 blockade in both the Th1 and Th2 conditions, CD4^+^ T cells produced a low amount of IL-2, which appeared to be sufficient for Th2 differentiation (**Extended Data Fig. 7i,j**).

Collectively, blockade of CD80/CD86 on APCs by sCTLA-4, and similarly by anti- CD80/CD86 antibodies, under TCR stimulation not only hinders proliferation of stimulated T cells but also selectively hampers their spontaneous Th1 differentiation in a sCTLA-4 dose-dependent manner, while allowing Th2 differentiation in the presence of IL-4.

### sCTLA-4 suppresses intestinal inflammation and tumor immunity, and facilitates wound healing

We next assessed immuno-suppressive and -modulatory functions of sCTLA-4 in mouse disease models. With the finding that a major source of sCTLA-4 was effector Tregs, we first examined *in vivo* suppressive function of Treg-derived sCTLA-4 to control colitis induced by cell transfer of WT CD45RB^hi^ naive CD4^+^ T cells into SCID mice^46^. Co-transfer of either S^+^M^-^ or S^-^M^+^ Tregs protected the mice from losing body weight and developing histologically evident colitis, with inhibition of colitogenic Th1/Th17 cells expansion, as effectively as co-transfer of WT S^+^M^+^ Tregs, whereas co-transfer of S^-^M^-^ Tregs did not (**Fig. 7a,b** and **Extended Data Fig. 8a**). Similar proportions of Tregs persisted in the SCID mice 8 weeks after each cell transfer; and adoptively transferred S^+^M^-^ or S^-^M^-^ Foxp3^+^ Tregs were comparable before transfer in the expression of various Treg signature molecules, such as CD25 and GITR, and in the transcription of immunosuppressive cytokines such as TGF-β, IL-10, and Ebi3 (IL-35 subunit) (**Extended Data Fig. 8b-d**). The results suggested a specific role of sCTLA-4 in the suppression of colitis.

**Fig. 7:**
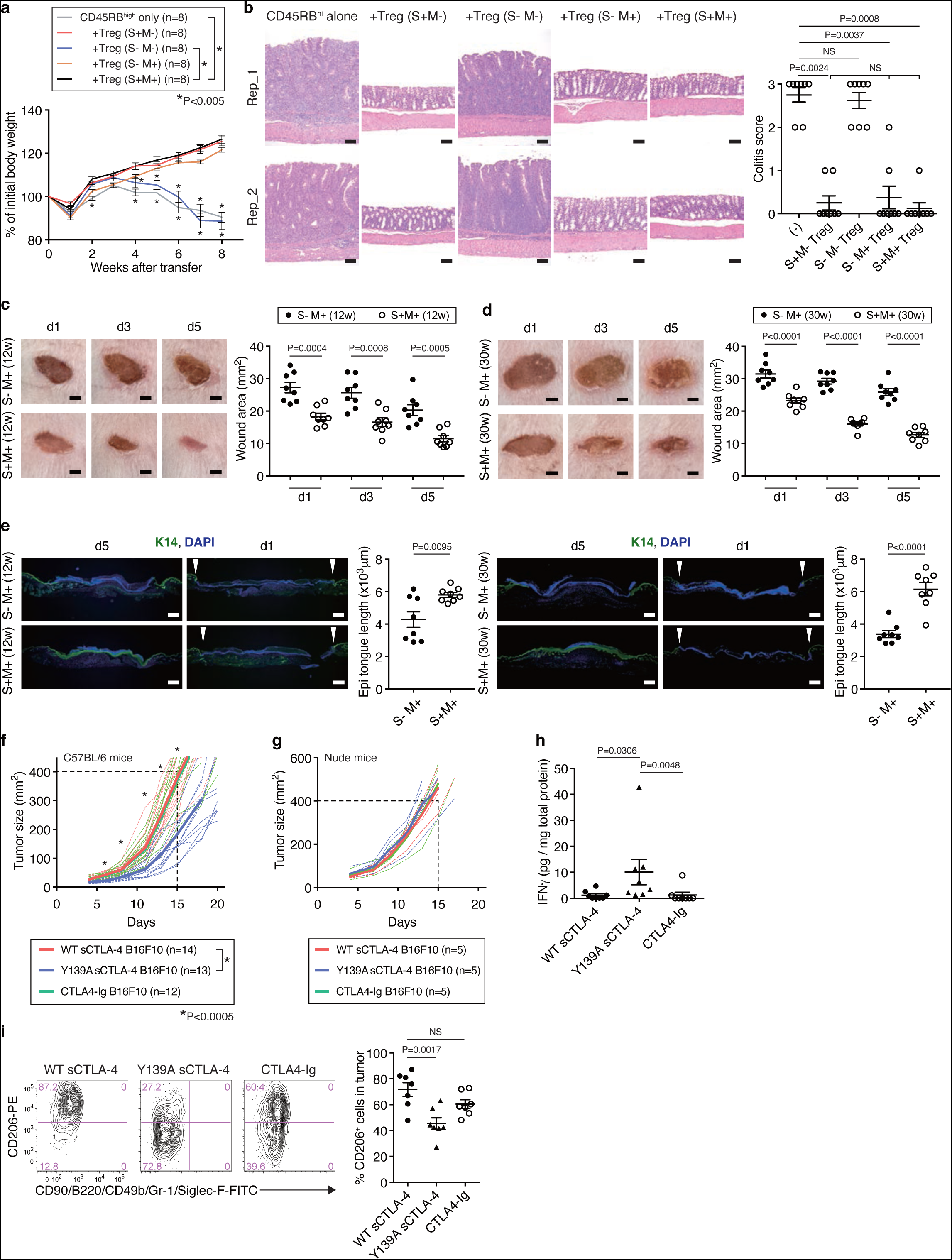
sCTLA-4 suppresses intestinal inflammation and tumor immunity, and facilitates wound healing. **a,** Prevention of colitis by S^+^M^-^ and S^-^M^+^ Tregs. Five-week-old C.B-17 SCID mice received intravenous injection of 4x105 Thy1.1^+^CD4^+^CD25^-^CD45RB^hi^Foxp3^-^(GFP^-^) T cells purified from eFOX Thy1.1^+^(CD90.1^+^) mice either alone or together with 3x105 Thy1.2^+^(CD90.2^+^) CD4^+^Foxp3^+^(GFP^+^) Tregs purified from 3-week-old FDG S^+^M^-^, FDG S^-^M^-^, FDG S^+^M^+^, or FDG S^-^M^+^ mice. Body weights are shown as percentages of initial body weights. Two-way ANOVA with Holm-Sidak multiple comparisons test. **b,** Histology of colitis and histological scores with sections from the colons of C.B-17 SCID mice described in (**a**). Scale bars represent 100 μm. The data were accessed by Kruskal-Wallis test followed by Dunn’s test. **c,d,** Skin lesions at the indicated time-points on the back of S^-^M^+^ and S^+^M^+^ mice at the age of 12 weeks (**c**) or at the age of 30 weeks (**d**). Skin wound was made by 6-mm biopsy punch. Average of duplicate wound area (mm2) per mouse was measured. Scale bars represent 2 mm. **e,** Re- epithelialization of wounded sites in S^-^M^+^ and S^+^M^+^ mice 5 days post wounding as described in (**c**) and (**d**). Sections are immunostained for K14^+^ basal epidermal keratinocytes (green) and co-stained with DAPI (blue). Re-epithelialization was quantified by the length of the tongue (Epi tongue length as illustrated in Extended Data Fig. 8g) of migrating K14^+^ epidermal keratinocytes. Scale bars represent 500 μm. Arrows show wound edges on day 1 with little signs of re-epithelialization. **f,g,** Inhibition of tumor immunity by sCTLA-4. Eight-week-old C57BL/6 mice (**f**) or nude mice (**g**) were subcutaneously inoculated with 1x106 B16F10 melanoma cells producing WT or Y139A sCTLA-4, or CTLA4-Ig. Length and width of tumor were measured every other day for 3 weeks to calculate tumor size (mm2). Lines shown in bold indicate average growth of individual tumors shown as dotted lines. **h,** Concentration of IFNγ in cell lysates of tumors 12 days after tumor cell inoculation. The data were accessed by Kruskal-Wallis test followed by Dunn’s test. **i,** Promotion of M2-like macrophage accumulation in tumors by tumor-secreted sCTLA-4. CD206 staining of tumor-infiltrated macrophages, defined as CD45^+^F4/80^hi^CD68^hi^ cells, in mice inoculated with indicated B16F10 melanoma cell lines and examined 12 days after tumor cell inoculation. All error bars denote the mean ± s.e.m. Data are representative or summary of at least two independent experiments with five or more mice (**a-i**). Statistical significances were determined by one-way ANOVA with Holm-Sidak multiple comparisons test in (**f,i**) or unpaired two-tailed *t*-test in (**c-e**).

We also investigated whether specific sCTLA-4 deficiency would affect tissue repair, for example, wound healing, which is hampered by chronic M1 polarization^47^. Assessment of the degree of wound healing on day 1, 3, and 5 after skin excision in 12- or 30-week-old S^-^M^+^ mice consistently showed a significant delay in wound closure compared with age-matched S^+^M^+^ mice (**Fig. 7c,d**). A comparison between 12- and 30-week-old S^-^M^+^ mice revealed that wound closure was significantly more delayed on day 5 in the latter (**Extended Data Fig. 8e**), which showed marked increases in Th1 and Tc1 cells, and M1-like macrophages (**Fig. 2g-k**). Moreover, S^-^M^+^ mice at both ages, when compared with age-matched S^+^M^+^ mice, showed significantly impaired re-epithelialization process assessed by measuring the length of epithelial tongue formation, which accompanied the migration of K14^+^ keratinocytes at the *stratum basale* of the epidermis underneath the wound (**Fig. 7e** and **Extended Data Fig. 8f,g**). In addition, in order to determine possible effects of sCTLA-4 on tumor immunity, we prepared B16F10 melanoma cell transfectants secreting WT or Y139A-mutated sCTLA-4, or CTLA4-Ig (**Extended Data Fig. 8h,i**). Each transduction did not affect *in vitro* proliferation of the tumor cells themselves (**Extended Data Fig. 8j**). When inoculated subcutaneously into C57BL/6 mice, WT sCTLA-4-secreting melanoma cells showed significantly faster tumor growth than Y139A sCTLA-4-secreting ones, while the two cell lines exhibited no difference in tumor growth in T-cell deficient athymic nude mice (**Fig. 7f,g**). In addition, compared with the tumor tissues of Y139A sCTLA-4-secreting melanoma, WT sCTLA-4- or CTLA4-Ig-secreting melanoma tissues showed significantly lower productions of IFNγ and significantly higher ratios of CD206^hi^ M2-like tumor-associated macrophages (**Fig. 7h,i**).

These results collectively indicate that sCTLA-4 produced by Tregs in particular is able to hamper type 1 immunity, while it is permissive to type 2 immunity, thereby suppressing chronic colon inflammation and facilitating tissue repair, but hindering tumor immunity.

## Discussion

A key finding in this study is that specific deficiency of sCTLA-4 evoked spontaneous activation of Th1 cells including self-reactive ones in otherwise normal mice and that sCTLA-4 production in CTLA-4 deficient autoimmune mice polarized activated Tconvs to Th2 cells. The finding indicates that sCTLA-4 possesses a distinct function in maintaining immunological tolerance and homeostasis by inhibiting functional polarization of activated Tconvs to Th1 cells but allowing their skewing to Th2 cells.

sCTLA-4 is immunosuppressive as demonstrated by the finding that S^+^M^-^ Tregs effectively suppressed experimental colitis and that sCTLA-4-specific deficiency induced much milder and less acute autoimmunity than mCTLA-4-specific deficiency as seen in S^-^M^+^ and S^+^M^-^ mice, respectively. On the other hand, the immunosuppressive function of mCTLA- 4 has been well documented in Tregs, as illustrated in our experiments by the effective suppression of colitis by S^-^M^+^ Tregs. Following physical cell contact between Tregs and APCs^48^, the CD80/CD86 molecules bound to mCTLA-4 on Tregs are transferred to the Treg cell surface by trogocytosis and endocytosed into Tregs by transendocytosis, leading to substantial reduction of CD80/CD86 expression on APCs and thus depriving Tconvs of co-stimulation/signal 2 needed for optimal activation^49, 50^. This Treg-mediated mCTLA-4- dependent CD80 down-regulation can also release PD-L1 from its cis-binding to CD80, thus granting free PD-L1 to engage and suppress PD-1^+^ effector Tconvs^50^. In contrast with these potent immuno-suppressive actions of mCTLA-4, sCTLA-4 in S^+^M^-^ mice did not down- regulate CD80 expression or increase free-PD-L1 on DCs. It is thus likely that Tregs supplement their mCTLA-4-dependent suppressive function by producing sCTLA-4 to further dampen any residual CD80/CD86 co-stimulation from Treg-conjugated APCs. sCTLA-4 may also be able to block CD80/CD86 expressed on neighboring Treg-nonconjugated APCs in a bystander manner, thereby suppressing activation and Th1 differentiation of Tconvs stimulated by such APCs (**Extended Data Fig. 9a)**. In Treg-conjugated APCs, it remains to be examined whether Treg-derived sCTLA-4 competes with Treg-expressed mCTLA-4 for binding to CD80/CD86 on an APC in a dose-dependent manner, thereby finely tuning the intensity of co-stimulatory and co-inhibitory signals delivered to Tconvs.

In addition to the immunosuppressive activity, sCTLA-4 is able to modulate Th1/Th2 differentiation in immune responses. To illustrate, S^-^M^+^ mice exhibited the activation/expansion of Th1, Th17, Tfh, and Tc1 cells, while S^+^M^-^ mice showed spontaneous and selective activation and expansion of Th2 cells. This is consistent with our *in vitro* results that anti-CD3 mAb stimulation of Th0 cells drove them to differentiate into Th1 cells spontaneously without any supplementation of cytokines^44, 45^ and that CD80/CD86 blockade by recombinant sCTLA-4 suppressed the spontaneous Th1 differentiation while allowing IL-4- dependent Th2 differentiation. Thus, depending on the number and the activation status of sCTLA-4-producing Treg and Tconv cell populations as well as their kinetics and intensity of its production, sCTLA-4 exerts distinct effects on the initiation and progression of immune responses. Namely, assuming that APCs are physiologically presenting self-antigens and microbial antigens from commensal microbes, CD80/CD86 blockade by sCTLA-4 especially produced by tissues Tregs, which are physiologically in an activated state^16^, may locally raise the threshold for activation of naïve Tconvs, including self-reactive T cells, in the non-inflammatory steady state (**Extended Data Fig. 9b-i**). A brief drop in Treg production of sCTLA-4 upon antigenic stimulation, may locally allow naïve Tconvs to become activated and initiate an immune response before a new wave of sCTLA-4 produced by activated Tregs, actively regulates the response (**Extended Data Fig. 9b-ii**). Once the immune response is in progress, abundant sCTLA-4 mainly produced by activated effector Tregs, together with sCTLA-4 from activated Tconvs, is able to inhibit the differentiation of antigen-stimulated Th0 cells towards Th1 cells, while skewing the differentiation towards Th2 cells, leading to resolution of inflammation and facilitation of tissue repair, as discussed below (**Extended Data Fig. 9b-iii**). Our results also indicate that sCTLA-4 is able to inhibit Tfh and Th17 differentiation as well, which needs further detailed mechanistic investigation.

There is accumulating evidence that TCR signal strength/duration, antigen dosage, and CD28 co-stimulatory signal intensity are key factors for determining Th1/Th2 differentiation^51–56^. It has been shown that CD80/CD86 blockade by CTLA-4-Ig suppress both Th1 and Th2 differentiation, which can be restored by increasing IL-2 availability^57^ and that CD80/CD86- deficient APCs strongly inhibit T cell production of IFNγ but permit IL-4 production^58, 59^. Thus, although both Th1 and Th2 differentiation depends on CD80/CD86 co-stimulation and IL-2, Th1 differentiation appears to require stronger and continuous TCR/CD28 signal than Th2 differentiation^56^. This is conceivable because TCR/CD28 stimulation is necessary for T-cell production of IL-2, a key factor for T-cell expression of IL-12Rβ2, and because T cell expression of CD40L is necessary for stimulating APCs via CD40 to produce IL-12^44, 45, 56^. Weak TCR signaling, on the other hand, induces Th0 cells to express GATA-3 and produce IL- 4 in an IL-4/STAT6-independent manner in an early phase of cell fate decision^60^. It is vital to note here that Th0 cells can initiate expression of IL-4 without pre-exposure to IL-4 and that the engagement of CD28 by CD80/86 is redundant for priming IL-4 production when IL-2 is available^57^. Moreover, IL-4 is known to facilitate differentiation of DC precursors towards cDC2 cells^61^, which may promote Th2 differentiation, as observed in S^+^M^-^ mice. The Th2 cells thus generated may then counterbalance Th1 and Th17 responses further via IL-4^62–64^. In addition, once Th2 cells are produced, they survive longer than Th1 cells, which may contribute to the accumulation of Th2 cells in sCTLA-4-abundant chronic inflammation^65^. Taken together, CD80/86 co-stimulation is a key element for determining the Th1/Th2 cell fate of antigen-activated naïve Tconvs, and the availability of sCTLA-4 is pivotal in this process. Thus, in autoimmune disease patients, high amounts of serum sCTLA-4, which were equivalent or higher than in autoimmune S^+^M^-^ mice, may be more than a mere clinical marker of chronic inflammation^12–15^. It may reflect active control of type 1 inflammation by sCTLA-4 locally produced by activated Tregs and Tconvs. In addition, some genetic polymorphisms at the *CTLA4* locus^6^, including the one with a loss-of-function effect on sCTLA-4 production in type 1 diabetes^5^, may affect the Th1-innhibitory effect of sCTLA-4, thereby contributing to autoimmune disease development in a similar manner as in S^-^M^+^ mice.

Th2 cells induced by sCTLA-4 produce various cytokines that activate/expand other immune cells and modify their phenotype and function. For example, *in vitro* IL-4/IL-10 treatment of macrophages isolated from S^-^M^-^ mice down-regulated their CD80 expression as observed *in vivo* with the macrophages in S^+^M^-^ mice. IL-4, IL-5, and IL-10 produced by Th2 cells in S^+^M^-^ mice drove the differentiation of macrophages into M2 type, and also activated and expanded IL-4^hi^ eosinophils, which reportedly contribute to tissue repair^66, 67^. Consistently, S^+^M^-^ mice showed an increase in Th2 and eosinophils in the blood and the lung, with upregulation of the serum CCL11, CCL22, and CCL24 (capable of recruiting Tregs, Th2 cells, and eosinophils), which are up-regulated by Th2 cytokines and downregulated by Th1 cytokines^28–32^. Type 2 immunity mediated by Th2 cytokines has been reported to be preventive of autoimmunity^68–75^, graft-versus-host disease^76^, and allograft rejection^77–79^. In addition to Th2 control of Th1, the preventive effects can be in part attributed to enhanced function of Tregs by Th2 cytokines. For example, IL-4 promoted suppressive and proliferative activities of Tregs and prolonged their survival by inhibiting apoptosis, thus enabling IL-4-treated alloantigen- primed Tregs to potently suppress graft rejection^76, 80–83^. IL-5, another prominent Th2 cytokine, enhanced the expansion of IL-5Rα^+^ antigen-specific Tregs to ameliorate experimental autoimmune neuritis^73^. Tregs can also produce IL-10^84, 85^, IL-13^86^, amphiregulin^87^, and CCN3^88^ to promote tissue repair as shown with various mouse models of tissue inflammation. Hence, these findings collectively suggest that sCTLA-4 produced by effector Tregs and effector/memory Tconvs under chronic inflammation amplifies type 2 immunity via Th2 cytokines, helps resolve inflammation, and facilitates tissue repair by inducing M2-like macrophages, activating IL-4^hi^ eosinophils, and enhancing Treg suppressive function.

The present study has demonstrated that sCTLA-4 is a good therapeutic candidate for immune regulation in various physiological and pathological settings. sCTLA-4 as a natural constituent of the immune system is less potent than engineered CTLA4-Ig (abatacept) in immune suppression because the former (which is monomer) is weaker than the latter (dimer) in the avidity for CD80/CD86^43^. Nevertheless, sCTLA-4 can be used to selectively suppress type 1 immunity and enhance type 2 immunity and to augment Treg-mediated suppression and tissue repair for treating chronic inflammatory diseases mediated by type 1 immunity. In tumor immunity setting, sCTLA-4 suppresses T cell production of IFNγ and facilitates accumulation of CD206^+^ M2-like tumor-associated macrophages in tumor microenvironment, resulting in attenuation of T-cell dependent anti-tumor immune responses. The serum sCTLA-4 level is reportedly correlated with the frequency of lung metastasis in a mouse melanoma model^10^. In addition, sCTLA-4-secreting effector Tregs are abundant in tumor tissues, hindering effective tumor immunity in humans and mice^89^. It is thus likely in cancer immunotherapy with the anti-CTLA-4 antibody ipilimumab that the antibody not only blocks mCTLA-4 on effector T cells and depletes mCTLA-4-expressing Tregs by antibody-dependent cellular cytotoxicity^7, 90, 91^, but also neutralizes sCTLA-4, thereby synergistically promoting Th1/Tc1-mediated anti-tumor immune responses. Though effective against tumor, the neutralization of sCTLA-4 by ipilimumab could also bring about higher risks of Th1-mediated autoimmunity as an adverse effect. In this regard, the serum concentration of sCTLA-4 could be a useful marker for assessing the mode and intensity of tumor immunity (and autoimmunity), especially the possible contribution of Tregs, in ipilimumab cancer immunotherapy.

In conclusion, sCTLA-4, which is predominantly produced by Tregs in steady and inflammatory states, possesses local and systemic immuno-suppressive and -modulatory activity that is essential for maintaining immunological self-tolerance and homeostasis especially through balancing type 1 and presumably type 3 immune responses with type 2 responses.

## Methods

### Mice

Female 6-8-week-old BALB/cAJcl, BALB/cAJcl-nu/nu, C57BL/6JJcl, and 5-week-old C.B17/Icr-scid mice were purchased from CLEA Japan. CTLA-4 KO (S^-^M^-^, targeted disruption of exon2 and exon3), CD45.1, CD80/CD86 KO, FLPeR, DEREG, Foxp3^IRES-DTR-GFP^ knock-in (FDG), enhanced green fluorescent protein (EGFP)-Foxp3 fusion protein knock-in (eFOX), and Thy1.1 mice were previously described^22, 92–97^. sCTLA-4^+^mCTLA-4^-^ (S^+^M^-^) mice were generated by standard molecular procedures. Briefly, BALB/c-I ES cells (a gift from B. Ledermann^98^) were transfected with a modified CTLA-4 targeting vector (RP23-146J17) in which a splicing acceptor site in front of exon 3 of CTLA-4 locus (86 bp) was removed. The targeting vector was constructed using a phage-based *Escherichia coli* homologous recombination system^99^. sCTLA-4^-^mCTLA-4^+^ (S^-^M^+^) mice were generated by conjugating *Ctla4* exon3 and exon4 in the CTLA-4 BAC transgene (RP23-146J17); the mice possessing the modified BAC were bred with S^-^M^-^ mice. S^+^M^-^ and S^-^M^-^ mice were used at 3 weeks of age, and S^-^M^+^ mice were used at 30-32 weeks of age. Unless otherwise stated, all the strains were backcrossed to the BALB/c background more than 10 times. The mice were maintained under specific pathogen-free conditions and treated in accordance with the guidelines for animal welfare approved by the Immunology Frontier Research Center, Osaka university or the Institute for Life and Medical Sciences, Kyoto University.

### Antibodies

The following antibodies (m, mouse; h, human) were used for experiments, with clones, vendors, and catalog numbers: anti-mCD3e (145-2C11, BD, 553057), anti-mCD28 (37.51, BD, 553294), anti-mCD4 (RM4-5, BioLegend, 100548 or 100550, or BD, 553044 or 563747), anti- mCD8a (53-6.7, BioLegend, 100741 or 100759, or BD, 553029), anti-mCD25 (PC61, BD, 553866 or 552880), anti-mCD5 (53-7.3, BD, 553018), anti-mGITR (DTA-1, BD, 558139), anti-mFR4 (eBio12A5, eBioscience, 25-5445-82), anti-mTCRβ (H57-597, BioLegend, 109229), anti-TCR DO11.10 (KJ1-26, Invitrogen, 17-5808-80), anti-mCTLA-4 (UC10-4F10-11, BD, 553720), anti-mPD-1 (RPM1-30, BioLegend, 109110, or 29F.1A12, BioLegend, 135218), anti-mCD44 (IM7, eBioscience, 17-0441-83 or 11-0441-82, or BD, 564587), anti- mCD69 (H1.2F3, eBioscience, 12-0691-82, or BD, 560689), anti-mCD103 (M290, BD, 557494), anti-mCD62L (MEL-14, BD, 562910, 560516, or 553151, or BioLegend, 104445), anti-mCD45RB (16A, BD, 553100, or BioLegend, 103312), anti-mCD223 (C9B7W, BD, 552380), anti-mCD90.2 (53-2.1, BD, 553004, or BioLegend, 140310), anti-mCD80 (16-10A1, eBioscience, 12-0801-81, or BioLegend, 104710), anti-mCD86 (PO3.1, eBioscience, 12-0861- 82 or 16-0861-85), anti-mLy-6C (HK1.4, BioLegend, 128037), anti-mLy-6G (1A8, BioLegend, 127606 or 127633), anti-mGr-1 (RB6-8C5, eBioscience, 11-5931-82, or BD, 553124), anti-m/hTER119 (TER119, eBioscience, 13-5921-82), anti-mSiglec-F (E50-2440, BD, 552126 or 564514), anti-mCD41 (MWReg30, BD, 553848 or 562957), anti-mCD117 (c-Kit) (2B8, BioLegend, 105816), anti-mCD200R3 (Ba13, BioLegend, 142206), anti-mCD24 (M1/69, BioLegend, 101822), anti-mFcεRIα (MAR-1, BioLegend, 134304), anti-mCD49b (HMα2, BioLegend, 103503, or DX5, BD, 553856), anti-m/hCD11b (M1/70, BioLegend, 101236 or 101242), anti-mCD11c (N418, eBioscience, 11-0114-85, or BioLegend, 117314, or HL3, BD, 550261, 553802, or 553800), anti-mCX3CR1 (2A9-1, eBioscience, 12-6099-41), anti-mF4/80 (BM8, BioLegend, 123110, 123122, or 123106), anti-mCD206 (C068C2, BioLegend, 141705), anti-mCD163 (S15049I, BioLegend, 155318), anti-mCD369 (218820, BD, 749795), anti- mCD209a (MMD3, BioLegend, 833004), anti-mEGR2 (erongr2, eBioscience, 12-6691-82), anti-mCD68 (FA-11, BioLegend, 137017), anti-mCD64 (X54-5/7.1, BioLegend, 139304), anti- mCD71 (RI7217, BioLegend, 113812), anti-I-A/I-E (M5/114.15.2, BioLegend, 107630), anti- mCD45 (30-F11, BioLegend, 103140), anti-mCD45.1 (A20, BD, 560578), anti-mCD45.2 (104, BD, 563051), anti-mCD45R/B220 (RA3-6B2, BD, 553088 or 560472, or BioLegend, 103244), anti-mCD19 (1D3, BD, 553784), anti-mCXCR5 (L138D7, BioLegend, 145511 or 145516), anti-mBcl-6 (K112-91, BD, 561522), anti-mCD38 (90, eBioscience, 17-0381-82, or BioLegend, 102714), anti-h/mGL-7 (GL-7, eBioscience, 48-5902-82), anti-mIgD (11-26c.2a, BioLegend, 405704), anti-mIgM (ll/41, eBioscience, 25-5790-82), anti-mCD138 (281-2, BD, 558626 or 553714), anti-mCD16/32 (93, BioLegend, 101302, or 2.4G2, BD, 553142), anti-mIL-2 (JES6-5H4, eBioscience, 11-7021-82), anti-mIFNγ (XMG1.2, BD, 557735, or eBioscience, 12-7311- 82), anti-mIL-4 (11B11, BD, 557728, or BioLegend, 504119), anti-m/hIL-5 (TRFK5, BioLegend, 504304), anti-mIL-17A (TC11-18H10.1, BD, 559502, or BioLegend, 506912 or 506925), anti-mIL-10 (JES5-16E3, BioLegend, 505014), anti-mFoxp3 (FJK-16s, eBioscience, 17-5773-82, 53-5773-82, or 48-5773-82), anti-mGata-3 (TWAJ, eBioscience, 50-9966-42), anti-mT-bet (4B10, BD, 561265), anti-mHelios (22F6, BioLegend, 137216), anti-mEomes (Dan11mag, eBioscience, 12-4875-80), anti-mNur77 (Nr4a1) (12.14, Invitrogen, 12-5965-82), Biotin Mouse Lineage Panel (BD, 559971), anti-m/hKeratin 14 (Poly9060, BioLegend, 906004), anti-hCD25 (M-A251, BD, 555432), anti-hCD4 (RPA-T4, BD, 561842), anti- hCD45RA (HI100, BD, 560675), anti-m/hKi-67 (B56, BD, 556026), anti-FLAG (M2, Sigma-Aldrich, F1804), anti-DYKDDDDK (L5, Invitrogen, MA1-142-A555), anti-m/hCTLA-4 (H- 126, Santa Cruz, sc-9094), anti-baculovirus gp64 (AcV1, eBioscience, 12-6991-82), FITC Streptavidin (BD, 554060), Brilliant Violet 421 Streptavidin (BioLegend, 405226), Ultra-LEAF Purified Armenian Hamster IgG Isotype Ctrl Antibody (HTK888, BioLegend, 400940), and Ultra-LEAF Purified Rat IgG2b, κ Isotype Ctrl Antibody (RTK4530, BioLegend, 400644). Anti-sCTLA-4-specific amino acids antibody (14K-12E) was generated by standard procedures for generating monoclonal antibody using sCTLA-4-unique peptides (NH_2_- AKEKKSSYNRGLCENAPNRARM-COOH) and biotinylated using the Biotin Labeling Kit - NH_2_ (Dojindo, LK03). For immunofluorescence staining, the antibodies described above were diluted in FACS buffer [1x HBSS (-) (nacalai tesque, 17460-15), 2% FBS (Gibco, 10437-028), and 0.01% Sodium azide (nacalai tesque, 31233-71)] at 1:200, except the anti- CD38/CXCR5/IgD antibodies at 1:100, the anti-Ki-67 antibody at 1:50, anti-human CD4/CD25/CD45RA antibodies at 1:10, and the Streptavidin FITC/Brilliant Violet 421 at 1:400. For macrophage/monocyte FACS staining, the True-Stain Monocyte Blocker (BioLegend, 426102) was used according to the manufacturer’s instructions. For MACS sorting, the CD11b MicroBeads (Miltenyi Biotec, 130-049-601), the CD11c MicroBeads (Miltenyi Biotec, 130-125-835), the CD90.2 (Thy1.2) MicroBeads (Miltenyi Biotec, 130-121-278), the Streptavidin MicroBeads (Miltenyi Biotec, 130-048-102), and the Anti-PE MicroBeads (Miltenyi Biotec, 130-048-801) were used according to the manufacturer’s instructions.

### Cell culture

Mouse primary T cells, splenocytes, lymphocytes, and blood cells were cultured in RPMI-1640 medium (nacalai tesque) supplemented with 10 mM HEPES solution (Sigma-Aldrich), 10% FBS (Gibco, 10437-028), 100 U/mL Penicillin-Streptomycin (Gibco), and 50 µM 2- mercaptoethanol (Sigma-Aldrich or Gibco). CD4^+^ T cells were activated by anti-CD3 mAb (0.5 μg/mL) and anti-CD28 mAb (1 μg/mL), anti-CD3/CD28 Dynabeads (Invitrogen), or anti-CD3 mAb (0.5 μg/mL) with T-cell depleted [CD90.2 depletion by the LD Columns (Miltenyi Biotec)] and Mytomycin C (KYOWA KIRIN)-treated (or non-treated) splenocytes in the presence of 100 U/mL human IL-2 (Imunase35, Shionogi). Splenocytes or lymphocytes prepared from 21-day-old S^+^M^-^, S^-^M^-^, and S^+^M^+^ mice were stimulated with 25 ng/mL PMA (Sigma-Aldrich) and 1 μM Ionomycin (Sigma-Aldrich) in the presence of Golgistop (Monensin, BD bioscience, 554724) for 5 hours. Cells were fixed and permeabilized by the Cytofix/Cytoperm and Perm/Wash (BD Biosciences, 554714) for staining cytokines, by the Foxp3/Transcription Factor Staining Buffer Set (eBioscience, 00-5523-00) for staining Foxp3, or by Transcription Factor Buffer Set (BD Biosciences, 562574) for co-staining of Foxp3 (transcription factors) and cytokines. Mouse primary macrophages were cultured in the presence of 10 ng/mL IL-4 (eBioscience, 14-8041-62) and 10 ng/mL IL-10 (eBioscience, 14- 8101-62) using the HydroCell 24 Multi-well plate (CellSeed). B16F10 melanoma cells were cultured in DMEM (nacalai tesque) supplemented with 10% FBS and Penicillin-Streptomycin. For baculovirus-insect cell expression system, Sf9 cells (Spodoptera frugiperda, Gibco, 11496015) were cultured using shake flask shaking at 130 rpm at 27°C in *Trichoplusia ni* Medium-Formulation Hink (TNM-FH) supplemented with 10% FBS, Penicillin-Streptomycin, and 0.1% Pluronic F-68 (Gibco); High Five (Hi5) cells (BTI-TN-5B1-4, Gibco, B85502) were cultured using shake flask shaking at 130 rpm at 27°C in Express Five SFM (Thermo Fisher Scientific) supplemented with Penicillin-Streptomycin, 10 U/mL heparin (MOCHIDA), and 18 mM L-glutamine (Gibco).

### ELISA

IFNγ, IL-4, TNF-α, and IL-6 were measured by the ELISA kit from eBioscience; IL-3, IL-10, and TSLP by the ELISA kit from Biolegend; CTLA-4, IL-33, and M-CSF by the ELISA kit from R&D Systems; Multiplex suspension array (IL-4, IL-5, IL-6, IL-13, IL-17A, and IFNγ) by the Bio-Plex Pro Mouse Cytokine Assay kit with the Bio-Plex 200 systems from Bio-Rad Laboratories; Serum antibodies by the Mouse Isotyping 6-plex Kit and the Mouse IgE FlowCytomix Simplex kit from eBioscience; Eotaxin (CCL11) and CCL22 by the LEGENDplex kit from Biolegend; Anti-mouse dsDNA IgG Ab and anti-mouse IgE by the ELISA kit from Sibayagi. Anti-parietal cell IgG Ab was determined as previously described^2^. Briefly, parietal-cell antigens were prepared from murine gastric corpus which was digested with 0.05 mg/mL Liberase DH (LIBTM-RO; Sigma-Aldrich) and 0.1 mg/mL DNase I (Roche); the proteins of the parietal-cell cytosol and membrane/organelle fractions were extracted with the ProteoExtract Subcellular Proteome Extraction Kit (Millipore) to exclude the nucleic and cytoskeletal fractions. Total protein from B16F10 melanoma was harvested using the M-PER mammalian protein extraction reagent (Thermo Fisher Scientific) with the Complete mini EDTA-free (Roche); the cell lysate was centrifuged for 10 minutes at 18,000 x g at 4°C and the supernatant was collected; concentration of whole proteins in the lysate was determined by the Pierce BCA Protein Assay Kit (Thermo Fisher Scientific) and concentration of IFNγ (pg/mg total protein) in the tumor lysate was quantified by ELISA. IL-10 production by peritoneal macrophages was measured by ELISA 24 hours after *in vitro* cell culture. For highly sensitive IL-2 quantification (0.064-5000 pg/mL), the IL-2 Mouse ProQuantum Immunoassay Kit (Invitrogen, A42892) was used.

### Quantitative Real-Time PCR (qPCR)

Total RNA was extracted by the RNeasy mini kit (QIAGEN). cDNA was synthesized by the SuperScript III or SuperScript IV VILO Master Mix (Invitrogen). Real-time PCR was performed by the StepOnePlus/QuantStudio3 Real-Time PCR System (Applied Biosystems) using the TaqMan Gene Expression Master Mix (Applied Biosystems) and the TaqMan Gene Expression Assays (FAM and VIC primer-limited) (Applied Biosystems) or performed by the LightCycler480 System (Roche) using the LightCycler480 Probes Master (Roche) and the Universal ProbeLibrary Probes (Roche). Transcripts were normalized to the mHprt/hHPRT transcript. Assay numbers are: Hprt (Hprt1), Mm01545399_m1; Il10, Mm00439614_m1; Ifng, Mm01168134_m1; Il4, Mm00445259_m1; Il17a, Mm00439618_m1; Il6, Mm00446190_m1; Ebi3, mm00469294_m1; Tgfb1, Mm03024053_m1; Nos2 (iNOS), Mm00440502_m1; Mrc1 (CD206), Mm00485148_m1; Retnla (Fizz1), Mm00445109_m1; Arg1, Mm00475988_m1; Tcf4, Mm00443210_m1; Irf7, Mm00516793_g1; Irf8, Mm00492567_m1; Batf3, Mm01318274_m1; Irf4, Mm00516431_m1; Klf4, Mm00516104_m1; Notch2, Mm00803077_m1; Zbtb46, Mm00511327_m1; Prdm1 (Blimp-1), Mm00476128_m1; CD80, Mm00711660_m1. The primer sequences of sCTLA-4 and mCTLA-4 are as follows: Mouse sCTLA-4; forward: 5’ - cgcagatttatgtcattgctaaag - 3’; reverse: 5’ - aaacggcctttcagttgatg - 3’; TaqMan probe (FAM NFQ-MGB): 5’ - aagaagtcctcttacaacagg - 3’. Mouse mCTLA-4; forward: 5’ - ggcaacgggacgcaga - 3’; reverse: 5’- cccaagctaactgcgacaagg - 3’; TaqMan probe (FAM NFQ-MGB): 5’- ttatgtcattgatccagaacc - 3’. Human sCTLA-4; forward: 5’ - ggacacgggactctacatctg - 3’; reverse: 5’ - aagagggcttcttttctttagca - 3’; Universal ProbeLibrary Probes #39 (Roche). Human mCTLA-4; forward: 5’ - ataggcaacggaacccagat - 3’; reverse: 5’ - cccgaactaactgctgcaa - 3’; Universal ProbeLibrary Probes #69 (Roche). Human HPRT1; forward: 5’ - gaccagtcaacaggggacat - 3’; reverse: 5’ - gtgtcaattatatcttccacaatcaag - 3’; Universal ProbeLibrary Probes #22 (Roche).

### Human samples

Blood samples were obtained from healthy adult volunteers (24-40 years old) with no overt symptomatic acute/chronic inflammatory or infectious diseases. This study was done according to the Helsinki declaration with process of obtaining informed consent and approval from the human ethics committee of the Institute for Life and Medical Sciences, Kyoto University. Isolation of CD4^+^CD45RA^+^CD25^+^ cells (naive Treg), CD4^+^CD45RA^-^ CD25^hi^ cells (effector Treg: < 1% of CD25^hi^), and CD4^+^CD45RA^-^CD25^int^ cells (Foxp3^+^ non-Treg) was conducted as previously described^17^.

### Plasmids

Mouse mCTLA-4 cDNA was cloned from BALB/c CD4^+^CD25^+^ T cells. To generate sCTLA-4 cDNA, mCTLA-4 cDNA was sub-cloned into the pCMV-Tag4A vector (Agilent Technologies) and an exon3 deletion construct was made by the KOD Plus Mutagenesis Kit (TOYOBO). Y139A (139 tyrosine > alanine) mutant sCTLA-4 was constructed by introducing mutations at the MYPPPY motif (ATGTACCCACCGCCATAC > ATGTACCCACCGCCAGCC). For retroviral gene transduction, wild-type (WT) sCTLA-4, Y139A mutant sCTLA-4, and CTLA4-Ig (pME18S-mCTLA4/WT-hIgG, a gift from T. Saito, RIKEN Center for Integrative Medical Sciences, Kanagawa, Japan) were sub-cloned into the pMCsIg retroviral vector (a gift from T. Kitamura, The Institute of Medical Sciences, The University of Tokyo, Tokyo, Japan) to generate pMCsIg-WT_sCTLA-4, pMCsIg- Y139A_sCTLA-4, and pMCsIg-CTLA4-Ig. For generation of recombinant baculoviruses, the C-terminal FLAG-tagged WT sCTLA-4 and Y139A sCTLA-4 in pCMV-Tag4A were sub- cloned into the pFastBac vector (Thermo Fisher Scientific) to generate pFastBac-WT_sCTLA- 4-FLAG and pFastBac-Y139A_sCTLA-4-FLAG.

### Expression and purification of sCTLA-4

Recombinant baculovirus was generated using the Bac-to-Bac Baculovirus Expression System (Thermo Fisher Scientific). High Five (Hi5) cells (BTI-TN-5B1-4) were infected with the baculovirus to produce C-terminal FLAG-tagged WT sCTLA-4 and Y139A sCTLA-4. The FLAG-tagged WT sCTLA-4 or Y139A sCTLA-4 in the culture supernatant was purified with the anti-FLAG M2 affinity gel (Sigma-Aldrich), washed with wash buffer (Phosphate buffered saline, 0.05% Tween-20), eluted by elution buffer [Phosphate buffered saline, 160 μg/mL 3xFLAG peptide (Sigma-Aldrich)], and dialyzed by PBS using the Slide-A-Lyzer G2 Dialysis Cassettes (10K MWCO) (Thermo Fisher Scientific). Concentration of sCTLA-4 was estimated by ELISA (R&D Systems, DY476) or SDS-PAGE/CBB staining with bovine serum albumin (BSA) standards.

### sCTLA-4- or CTLA4-Ig-producing B16F10 melanoma

Gene transduction of retroviral constructs described above (pMCsIg-WT_sCTLA-4, pMCsIg- Y139A_sCTLA-4 and pMCsIg-CTLA4-Ig) was conducted as previously described^100^. Although B16F10 cells are adherent cells, transduction was performed using a spinoculation protocol with 8 μg/mL Polybrene (Millipore) for 90 minutes at 1,000 x g at 32°C. After the retroviral transduction, GFP^+^ transduced cells were sorted and enriched with the BD FACSAria II twice. The expressions of the sCTLA-4 and CTLA4-Ig were assessed by FACS and ELISA.

### *In vitro* proliferation assay and Th1/Th2 polarization assay with sCTLA-4

For *in vitro* proliferation assay assessed by CSFE dye dilution, CD4^+^CD25^-^CD62L^hi^ naïve T cells from CD45.1 mice were labeled with 1 μM CFSE (Thermo Fisher Scientific), and the labeled T cells were used as responder T cells. In the presence of 0.5-1 x 104 Mitomycin C (KYOWA KIRIN)-treated CD45.2^+^ antigen presenting cells (T-cell depleted splenocytes) and soluble anti-CD3 mAb (0.5 μg/mL), 5 x 104 responder T cells were cultured with 1 μg/mL WT sCTLA-4, Y139A sCTLA-4, or CTLA4-Ig for 86-90 hours. For *in vitro* Th1/Th2 polarization assay, 5 x 104 CD4^+^CD25^-^CD62L^hi^ cells (CD45.2^+^) were stimulated with 0.5 μg/mL anti-CD3 mAb and 1 x 105 T-cell depleted splenocytes (CD45.1^+^) for 0-120 hours with 0.1-5.0 μg/mL WT or Y139A sCTLA-4, CTLA4-Ig, or 10 μg/mL anti-CD80 mAb (16-10A1, BioLegend, 104748) or anti-CD86 mAb (PO3.1, eBioscience, 16-0861-85), and then stimulated with PMA/Ionomycin/Monensin for 5 hours. The Ultra-LEAF Purified Armenian Hamster IgG Isotype Ctrl Antibody (BioLegend, 400940) or the Ultra-LEAF Purified Rat IgG2b, κ Isotype Ctrl Antibody (BioLegend, 400644) were used as isotype controls (10 μg/mL) of the anti-CD80 or CD86 mAb, respectively. Cells were fixed and permeabilized by the Transcription Factor Buffer Set (BD, 562574) for co-staining of transcription factors and cytokines. Th1 (minimal) condition, 10 μg/mL anti-IL-4 mAb (11B11, BioLegend, 504122); Th2 condition, 10 ng/mL IL-4 [eBioscience, 14-8041-62 (discontinued), or PeproTech, 214-14]. The 96-well round (U) bottom plate (Thermo Scientific, 163320) was used for T-cell culture.

### Immunofluorescence

CD11c^hi^ splenocytes which were stimulated by 1 μg/mL LPS (Sigma-Aldrich) for 24 hours were smeared onto MAS-coated slides (MATUNAMI GLASS) using the Smear Gell kit (NIPPON Genetics). Stained samples were observed with the FluoView FV10i confocal microscope (Olympus Life Science).

### Immunoprecipitation and immunoblot analysis

Immunoprecipitation (IP) was performed in binding buffer (50 mM Hepes, pH 7.5, 150 mM NaCl, 5% glycerol, 1% BSA, 0.05% Tween-20) containing 1 μg FLAG-tagged sCTLA-4 and 1 μg mCD80 Fc (human IgG1) chimera (R&D, 740-B1), 1 μg mCD86 Fc (human IgG1) chimera (R&D, 741-B2), or 1 μg Human IgG1 Fc (R&D, 110-HG) in the presence or absence of 1 μg mCD152/Fc (CTLA4-Ig, mouse IgG2a, non-cytolytic) chimera (Sigma-Aldrich, C4358). The anti-human IgG Fc Antibody (BioLegend) and the Sheep Anti-Rat IgG Dynabeads (Thermo Fisher Scientific, 11035) were used for pull-down assay. The pull-downed proteins were separated by SDS-PAGE on 10-to-20% gradient gel (ATTO) and the recombinant FLAG-tagged WT sCTLA-4 was detected by anti-FLAG M2 antibody (Sigma-Aldrich).

### Immunization of DO11.10 mice with ovalbumin

Eight- to 12-week-old DEREG (or FDG) DO11.10 TCR transgenic mice subcutaneously received 50 μL injections of a 1:1 emulsion of adjuvant complete Freund H37 Ra (BD Difco, 231131) and 2 mg/mL grade V ovalbumin (Sigma-Aldrich, A5503) in four points of their back near axillary/brachial lymph nodes or inguinal lymph nodes. On day 0, 7, 14, and 21, CD4^+^Foxp3^+^(GFP^+^) Tregs (purity > 99%) or CD4^+^Foxp3^-^(GFP^-^)DO11.10^+^ Tconvs (purity > 99%) were sorted from the draining swollen lymph nodes for quantification of sCTLA-4 and mCTLA-4 mRNA by qPCR.

### Cell transfer and tumor inoculation

Five-week-old C.B17/Icr-scid mice intravenously received 4 x 105 CD4^+^CD25^-^CD45RB^hi^ Foxp3^-^(GFP^-^) T cells purified from spleens and lymph nodes of eFOX Thy1.1^+^(CD90.1^+^) congenic mice either alone or together with 3 x 105 CD4^+^Foxp3^+^(GFP^+^) T cells from 3-week-old FDG S^+^M^-^, FDG S^-^M^-^, FDG S^+^M^+^, or FDG S^-^M^+^ mice. For tumor-bearing mouse models, 1 x 106 B16F10 melanocytes were subcutaneously inoculated into C57BL/6 or BALB/cAJcl-nu/nu mice. The experiments were terminated when a solid tumor reached criteria for endpoint; tumor size > 400 mm2 (20 mm x 20 mm).

### Isolation of lung cells, splenocytes, blood cells, and lamina propria cells

For single-cell preparation of immunocytes in the lung, lung tissues were perfused to remove the blood, homogenized and digested in 2.5 mL of the enzyme solution [1 mg/mL Hyaluronidase Type IV-S (Sigma-Aldrich), 0.05 mg/mL Liberase TM (LIBTM-RO; Sigma-Aldrich) and 0.1 mg/mL DNase I (Roche; Sigma-Aldrich)] with the program “m_lung_01.01 C Tube” of the gentleMACS Dissociator (Miltenyi Biotec) in the Bioshaker (TAITEC, BR- 22FP) at 220 rpm for 30 minutes at 37°C, and further homogenized with the program “m_lung_02.01 C Tube”. The cell suspensions thus prepared were passed through the 40-μm Cell Strainers (Falcon; Corning), stained with antibodies, and analyzed by FACS or sorted for Siglec-F^+^CD11c^+^CD11b^int^CD64^+^ alveolar macrophages by BD FACSAria II. For single-cell preparation of splenic CD11c^+^ dendritic cells, perfused and chopped spleen tissues were treated with 2 mL of the enzyme solution [0.05 mg/mL Liberase TM (LIBTM-RO; Sigma-Aldrich) and 0.1 mg/mL DNase I (Roche; Sigma-Aldrich)] in the Bioshaker (TAITEC, BR-22FP) at 220 rpm for 30 minutes at 37°C, and homogenized by straining through the 40-μm Cell Strainers (Falcon; Corning) with syringe plungers. CD11c^+^ dendritic cells in the cell suspensions were enriched by the CD11c MicroBeads (Miltenyi Biotec) and MACS Cell Separation Columns (Miltenyi Biotec), and sorted by the BD FACSAria II. For removal of erythrocytes from blood samples, heparinized mouse whole blood [Heparin Sodium (MOCHIDA, 224122458)] in the borosilicate glass tube (Fisher Scientific, 14-961-30) was mixed with an equal volume of 2% dextran solution [2% Dextran T500 (Pharmacosmos, 5510-0500-9006, Mw 450,000-550,000), 0.02% Sodium azide (nacalai tesque) in 1xPBS] and kept at 37°C for 30 minutes to precipitate erythrocytes. The supernatant was collected and treated with the Red Blood Cell Lysing Buffer (Sigma-Aldrich, R7757) to completely remove erythrocytes, and used for FACS analysis. Large intestinal lamina propria lymphocytes and myeloid cells were isolated as previously described^101^. After digesting the colonic mucosa, CD45^+^Siglec-F^-^Ly6G^-^CD11b^+^F4/80^+^ colonic macrophages were sorted by the BD FACSAria II for qPCR.

### Histological Analysis

Histological analysis of colon was conducted as previously described^46, 97^. Briefly, hematoxylin and eosin (HE) stained sections were histologically scored in a double-blinded manner, based on the following criteria. Colitis score: 0 (normal colon), small number of leukocytes in mucosa and large number of goblet cells in crypts; 1, slight epithelial cell hyperplasia and increased number of leukocytes in mucosa; 2, pronounced epithelial cell hyperplasia, significant leukocytic infiltrate in mucosa and submucosa, and depletion of mucin-secreting goblet cells; 3, marked epithelial cell hyperplasia, extensive leukocytic infiltrate in mucosa, submucosa, and *tunica muscularis*, ulceration, and significant depletion of mucin-secreting goblet cells. HE stain of the lung sections were assessed with the BZ-X710 microscope (KEYENCE). The ratio of whole lung area and lymphocyte infiltrated area (cell foci: > 1,000 μm2) was calculated and quantified by the BZ-X Analyzer software v1.4.1.1 (KEYENCE).

### RNA seq

Total RNA was extracted from sorted CD11b^hi^F4/80^hi^ peritoneal macrophages using the miRNeasy Mini Kit (QIAGEN). Purity of sorted macrophages were > 99%. cDNA libraries prepared by the TruSeq RNA Sample Prep Kit (Illumina) were sequenced using the HiSeq 2000 sequencing system (Illumina). Read sequences (paired-end) were mapped to the reference genome (mm10) with the TopHat (v2.0.13). Overall read mapping rates from all samples were > 94%. Expression levels and the statistical significance were calculated and tested with the Cuffdiff, which is included in the Cufflinks package (v2.2.1). The output files were visualized with the CummeRbund (v2.8.2) and gplots packages (v3.0.1) within the R environment (v3.1.2). For gene ontology (GO) enrichment analysis, read counts for each gene were calculated with the htseq-count which is the script based on the HTSeq (v0.7.1); a differential expression analysis was performed with the edgeR (v3.8.6) packages of the Bioconductor; the overrepresentation analysis (ORA) method with the ErmineJ software^102^ (v3.0.2) was used to assess differentially expressed genes. For mouse or human CTLA-4 exon3 skipping analysis, fastq files obtained from the EBI (European Bioinformatics Institute) database were trimmed and quality-controlled using the trim-galore (v0.6.7) to remove adaptor sequences and low-quality bases. The validated fastq reads were mapped to the genome (mouse, UCSC mm10; human, UCSC hg38) using the hisat2 (v2.2.0). Analysis of junction-spanning reads (output file: SE.MATS.JC.txt) using the sorted bam files with the rMATS (v4.1.2) was visualized by the sashimi-plot (rmats2sashimiplot, v2.0.2). Intron regions were compressed 5-fold. Mouse *Ctla4*, chr1: +:60912423:60915832; Human *CTLA4*, chr2: +:203870586:203873960. For KEGG (Kyoto Encyclopedia of Genes and Genomes) pathway analysis, an index was built for the following star-rsem analysis using GRCm39.genome.fa and gencode.vM26.annotation.gtf from GENCODE. Using the created index, fastaq reads were mapped to the genome using the star (v2.7.10a), and the resulting bam files were used for quantitative expression analysis using the rsem (v1.2.28). The obtained count matrix was used for KEGG pathway analysis using the iDEP (v0.96) and visualized by the Pathview^103–105^. For scRNA-seq analysis, filtered gene-barcode matrices were generated from fastq files obtained from the EBI database with refdata (GRCh38-2020-A) using the cellranger (v6.1.2, 10x Genomics). UMAP dimensionality reduction, clustering, and gene expression analysis were performed using the scanpy (v1.9.1) in the Python (v3.10.8) environment. Human breast cancer cell data were quality-controlled using the scanpy (v1.9.1) with the following conditions (min_cells = 3, min_genes = 200, n_genes_by_counts < 5000, pct_counts_mt < 15). After log-normalization (target_sum = 1e4), highly variable genes (min_mean = 0.0125, max_mean = 3, min_disp = 0.5, flavor = seurat) were selected for cell clustering analysis. For dimensionality reduction, PCA (n_comps = 50) was performed, and UMAP was computed using the neighbors parameter (n_neighbors = 10, n_pcs = 40). The Leiden algorithm (leidenalg, v0.9.0) was used for clustering.

### Blood chemistry and cell analysis

Blood amylase (AMY), AST, ALT, albumin (ALB), and total bilirubin (T-BIL) were determined by the 7180 Clinical Analyzer (Hitachi High-Tech) using the following reagents: L-Type Amylase (FUJIFILM Wako Chemicals); L-Type AST.J2 (FUJIFILM Wako Chemicals); L-Type ALT.J2 (FUJIFILM Wako Chemicals); Albumin II-HA (FUJIFILM Wako Chemicals); Nescort VL T-BIL (alfresa). The numbers of white blood cells (WBC) and red blood cells (RBC), and hematocrit (HCT, %) and hemoglobin (HGB, g/dL) were determined by the veterinary haematology analyser (Celltac alfa, MEK-6450, Nihon Kohden).

### Wound healing model and immunofluorescence of re-epithelialization

Excision of mouse back skin and wound healing quantification were performed as previously described^106^. Briefly, duplicate wounds were generated by the 6-mm biopsy punch (KAI medical) on the shaved back skin that disinfected with ethanol. The average wound area (mm2) of the duplicates was photographed with a scale and determined using the ImageJ software version 1.53a (National Institutes of Health) on day 1, 3, and 5 post wounding. Five days after wounding, back-skin tissue was collected for immunostaining, fixed in 4% paraformaldehyde in PBS at 4°C for 48 hours, embedded in paraffin, sectioned (5 μm) at longer axis of wound area, rehydrated, subjected to antigen retrieval with steam autoclaving at 121°C for 5 min in 10 mM sodium citrate (pH 6.0), and stained with anti-Keratin 14 antibody (Poly9060, Chicken polyclonal IgY, 1:500, BioLegend), followed by anti-Chicken IgY Cross-Adsorbed Secondary Antibody conjugated with Alexa Fluor Plus 488 (Goat polyclonal IgG, 1:1,000, Thermo Fisher Scientific) and DAPI (Thermo Fisher Scientific). Images of re-epithelialization underneath the wound were captured with the BZ-X710 microscope (KEYENCE) and the length of K14^+^ epidermal keratinocyte layers migrated from the wound edges was quantified by the ImageJ software as illustrated in Extended Data Figure 8g.

### Statistical analysis

In all the figures, each data point represents a measured value with a randomly selected animal, an independently conducted experiment, or mRNA level of a sample from different animals. Unless otherwise stated, data were analyzed by the one-way ANOVA followed by, if applicable, the Holm-Sidak test for multiple groups, or the unpaired two-tailed *t*-test for two groups using the GraphPad Prism version 7.0d (GraphPad software). For nonparametric test for multiple groups, the Kruskal-Wallis test followed by the Dunn’s multiple comparisons test was applied. When statistical analysis was conducted by the two-way ANOVA followed by the Holm-Sidak test for multiple groups, the non-parametric Mann-Whitney test for two groups, or the Log-rank test for comparing survival curves, it is stated in each figure legend. Error bars denote the mean ± standard error of the mean (s.e.m.) or standard deviation (s.d.), which are noted in each figure legend. *P*-values > 0.0500 were considered as NS (not significant). In cases where the *P*-value was 0.0500-0.1000 (NS), the value was described on figures.

## Data availability

The RNA-seq data reported in this study can be accessed at the DNA Data Bank of Japan (DDBJ) under accession number DRA013129. The data set has been mirrored to the NCBI Sequence Read Archive (SRA) and the European Bioinformatics Institute (EBI), and also available at the repositories. Previously published data were used for this work: bulk RNA-seq; GSE108184^107^, GSE120280^108^, GSE112341^109^, GSE89225^110^, GSE178132^111^, GSE106396^112^, GSE121836^113^, and GSE76138^114^: scRNA-seq; GSE176078^115^ from the Gene Expression Omnibus (GEO) database. Source data are provided with this paper.

## Acknowledgements

We are deeply grateful to all the members of Sakaguchi laboratory, especially to C. Tay for critical reading of the manuscript. We would like to thank Y. Esaki (NPO for Biotechnology Research and Development, Osaka university) for microinjection service of ES clones we established, H. Miyachi and S. Kitano (Reproductive Engineering Team, Institute for Life and Medical Sciences, Kyoto University) for the microinjection service of BAC vectors we constructed, B. Ledermann (Novartis Institutes for BioMedical Research) for kindly providing BALB/c-I ES cell line, and T. Saito (RIKEN Center for Integrative Medical Sciences) for providing pME18S-mCTLA4/WT-hIgG vector. This study was supported by Grants-in-Aid for Specially Promoted Research from Japan Society for the Promotion of Science under Grant Number 16H06295, and Japan Agency for Medical Research and Development (AMED) under Grant Number JP15gm0410016 (CREST) and JP18gm0010005 (LEAP).

## Contributions

M.O. and S.S. designed the experiments. M.O. performed all the experiments and analyzed the data; M.O. and S.S. wrote the manuscript.

## Abbreviations

CTLA-4: Cytotoxic T-Lymphocyte Antigen 4
sCTLA-4: Soluble CTLA-4
mCTLA-4: Membrane CTLA-4
DEREG mice: DEpletion of REGulatory T cells mice
FDG: Foxp3^IRES- DTR-GFP^ knock-in mice
eFOX: enhanced green fluorescent protein (eGFP)-Foxp3 fusion protein knock-in mice
BAC: Bacterial Artificial Chromosome
ILC: Innate Lymphoid Cell
cDC: classical/conventional Dendritic Cells
SCID: Severe Combined Immunodeficiency.

## Figure legends

**Extended Data Fig. 1:**
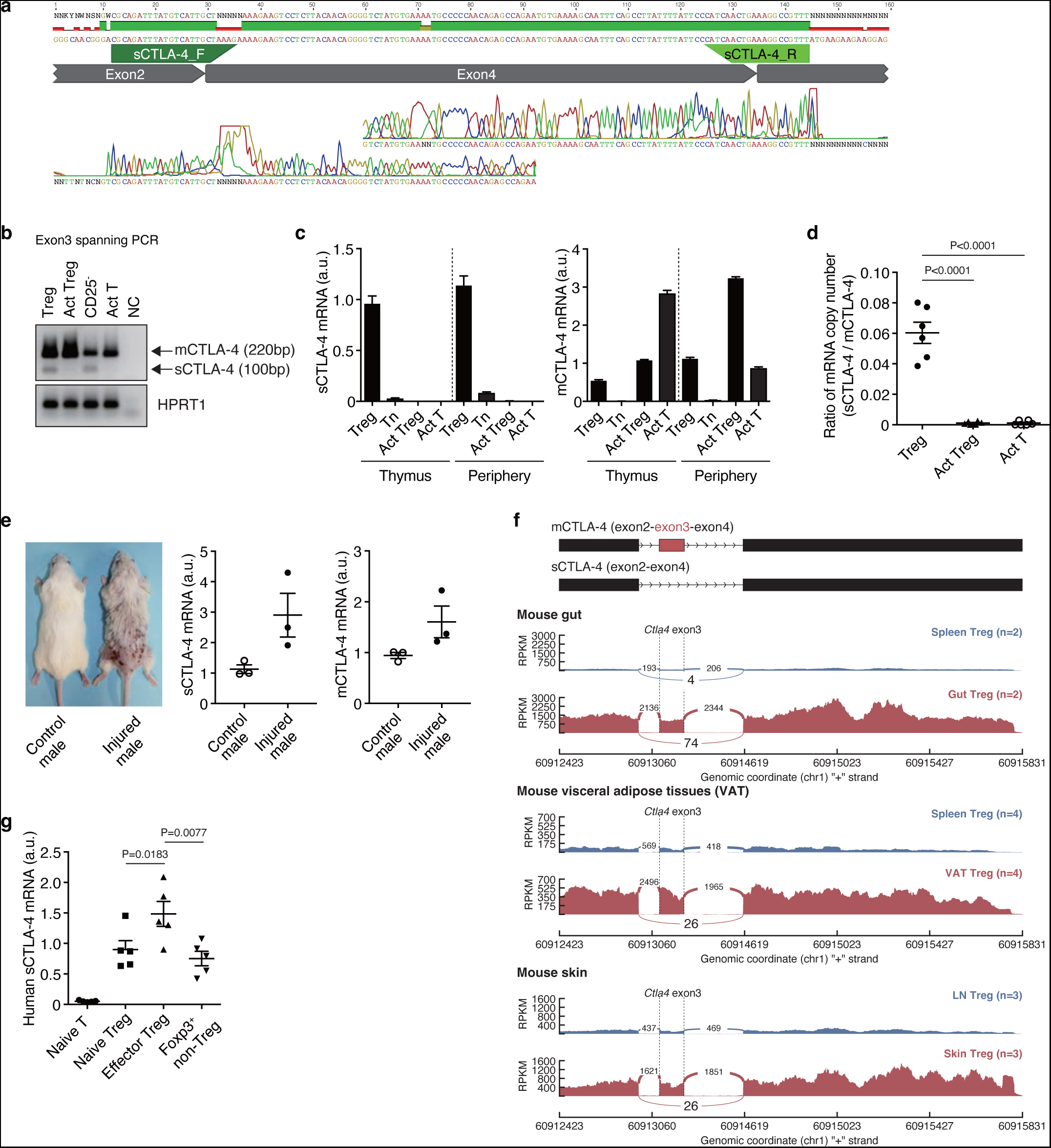
Predominant expression of sCTLA-4 by Tregs, especially tissue Tregs, in mice and humans. **a,** Evaluation of sCTLA-4 qPCR detection. Real-time PCR product amplified by sCTLA-4 detection primers (custom designed) were sequenced for evaluating accuracy. **b,** Specific diminishment of sCTLA-4 mRNA in CD4^+^ T cells after TCR stimulation. CTLA-4 cDNAs prepared from indicated T cells were amplified using exon3 spanning primers (The forward primer was designed based on exon2 sequence; the reverse primer was designed based on exon4 sequence.) and electrophoresed on 2.5% agarose gel. The agarose gel was stained with SYBR green. **c,** Relative mRNA expression of sCTLA-4 and mCTLA-4 in CD4^+^CD25^+^ cells (Treg), CD4^+^CD25^-^ cells (Tn), CD4^+^CD25^+^ cells with 3-day activation (act Treg), and CD4^+^CD25^-^ cells with 3-day activation (act T) from thymus or peripheral lymphoid organs. **d,** Ratio of sCTLA-4/mCTLA-4 mRNA copy number determined by real-time PCR absolute quantification. Treg, CD4^+^CD25^+^ cells; Act Treg, CD4^+^CD25^+^ cells with 3-day activation; Act T, CD4^+^CD25^-^ cells with 3-day activation using anti-CD3/CD28 mAb-coated Dynabeads. **e,** Upregulation of sCTLA-4 in Tregs isolated from chronic wounded mice. Relative mRNA expression of sCTLA-4 and mCTLA-4 in Foxp3^+^(GFP^+^)CD4^+^ T cells isolated from surface lymph nodes of DEREG male mice with ulcerated skin were compared with atraumatic mice with healthy skin. **f,** Visualization of CTLA-4 exon3 skipping (sCTLA-4 expression) by sashimi-plots. Tissue Tregs in the gut, visceral adipose tissue, and skin expressed relatively more exon3-skipped transcripts than Tregs in secondary lymphoid organs. Intron regions were compressed 5-fold. Mouse gut, GSE106396; Mouse visceral adipose tissues, GSE121836; Mouse skin, GSE76138 from the Gene Expression Omnibus (GEO) database. **g,** Upregulation of sCTLA-4 in human effector Tregs in peripheral blood. Relative mRNA expression of human sCTLA-4 in CD4^+^CD45RA^+^CD25^+^ cells (naive Treg), CD4^+^CD45RA^-^CD25^hi^ cells (effector Treg: < 1% of CD25^hi^), and CD4^+^CD45RA^-^CD25^int^ cells (Foxp3^+^ non-Treg) were determined. Error bars denote the mean ± s.e.m. in (**d-e,g**) or the mean ± s.d. in (**c**). Data are representative or summary of at least two independent experiments with two or more mice/samples (**a-e**) or with five human samples (**g**). Statistical significances were determined by one-way ANOVA with Holm-Sidak multiple comparisons test in (**d,g**) or unpaired two-tailed *t*-test in (**e**).

**Extended Data Fig. 2:**
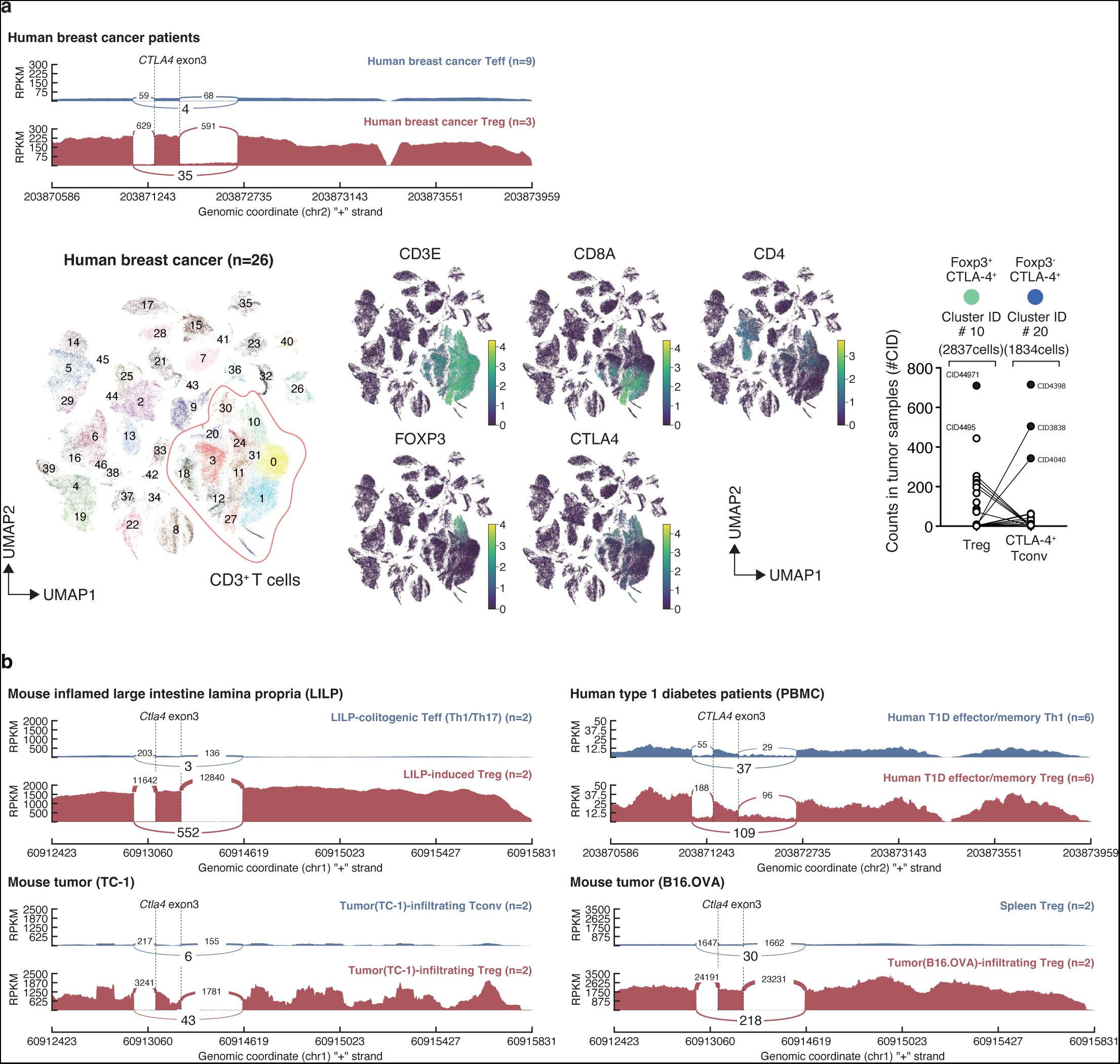
Predominant expression of sCTLA-4 by Tregs compared with effector Tconvs in tumor and inflammatory tissues in mice and humans. **a,** sCTLA-4 transcription by Tregs and effector Tconvs in human breast cancers. Visualization of CTLA-4 exon3 skipping (sCTLA-4 expression) by sashimi-plots. Intron regions were compressed 5-fold. The numbers of tumor-infiltrating Foxp3^+^CTLA-4^+^CD4^+^ Tregs (Cluster ID #10) and Foxp3^-^CTLA-4^+^CD4^+^ Tconvs (Cluster ID #20) in breast cancer tissues as assessed by single-cell RNA-seq are also shown. The patient sample IDs, CID44971, CID4040, CID3838, and CID4398, were outliers (closed circles) analyzed by ROUT method (Q = 1%), but were included in the data. **b,** sCTLA-4 transcription by Tregs and effector Tconvs in mouse colitis, mouse tumor, and human PBMCs of type 1 diabetes patients (or B16.OVA tumor-infiltrating Tregs vs. splenic Tregs). The data sets of bulk or single-cell RNA-seq used for the analyses are the following: GSE89225 (human breast cancers), GSE176078 (single-cell of human breast cancers), GSE108184 (mouse large intestine lamina propria with pathobiont-induced colitis), GSE120280 (mouse TC-1 tumor), GSE112341 (PBMCs of human type 1 diabetes patients), and GSE178132 (mouse B16.OVA tumor) from the Gene Expression Omnibus (GEO) database. PBMC, Peripheral blood mononuclear cell.

**Extended Data Fig. 3:**
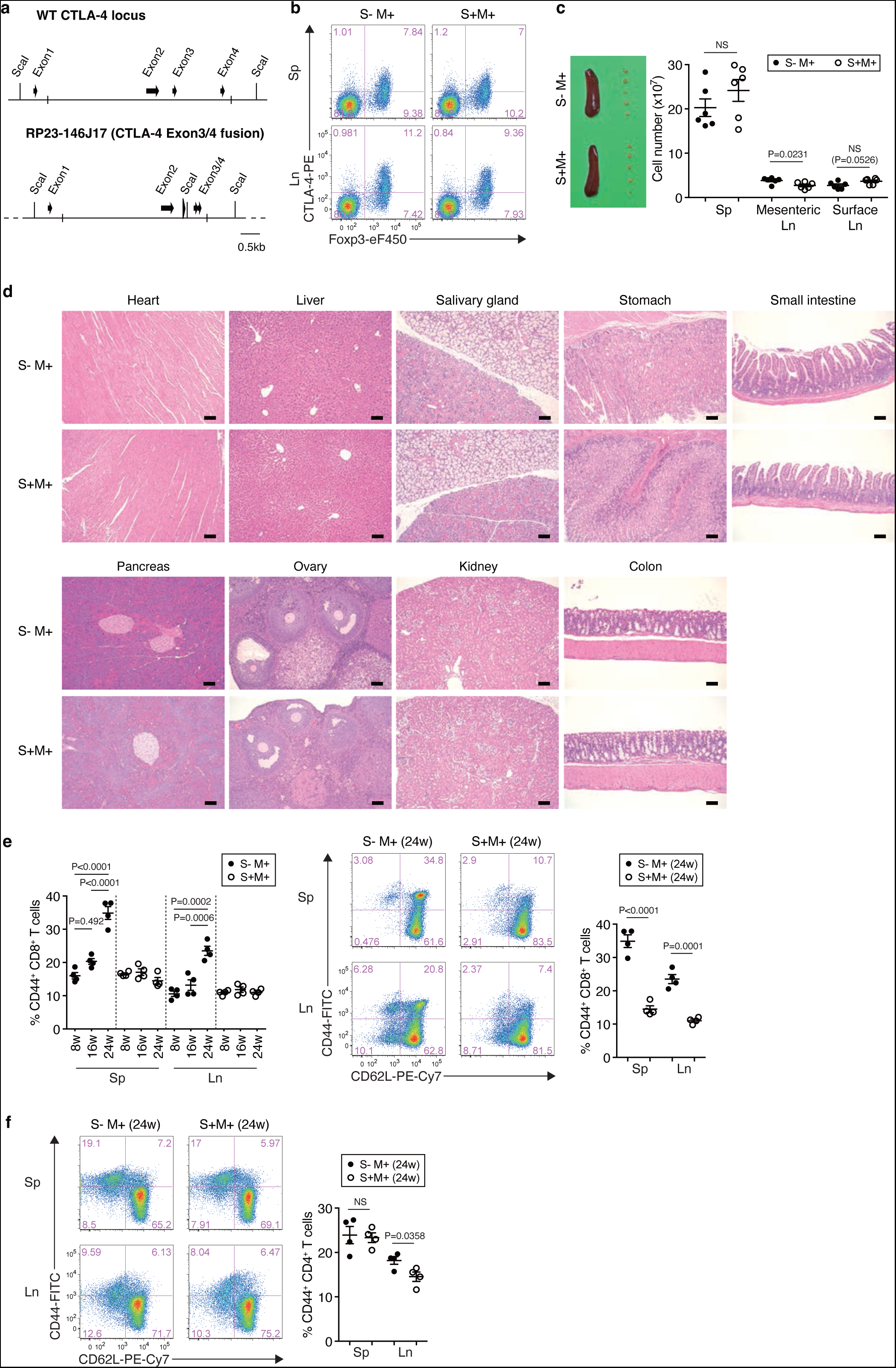
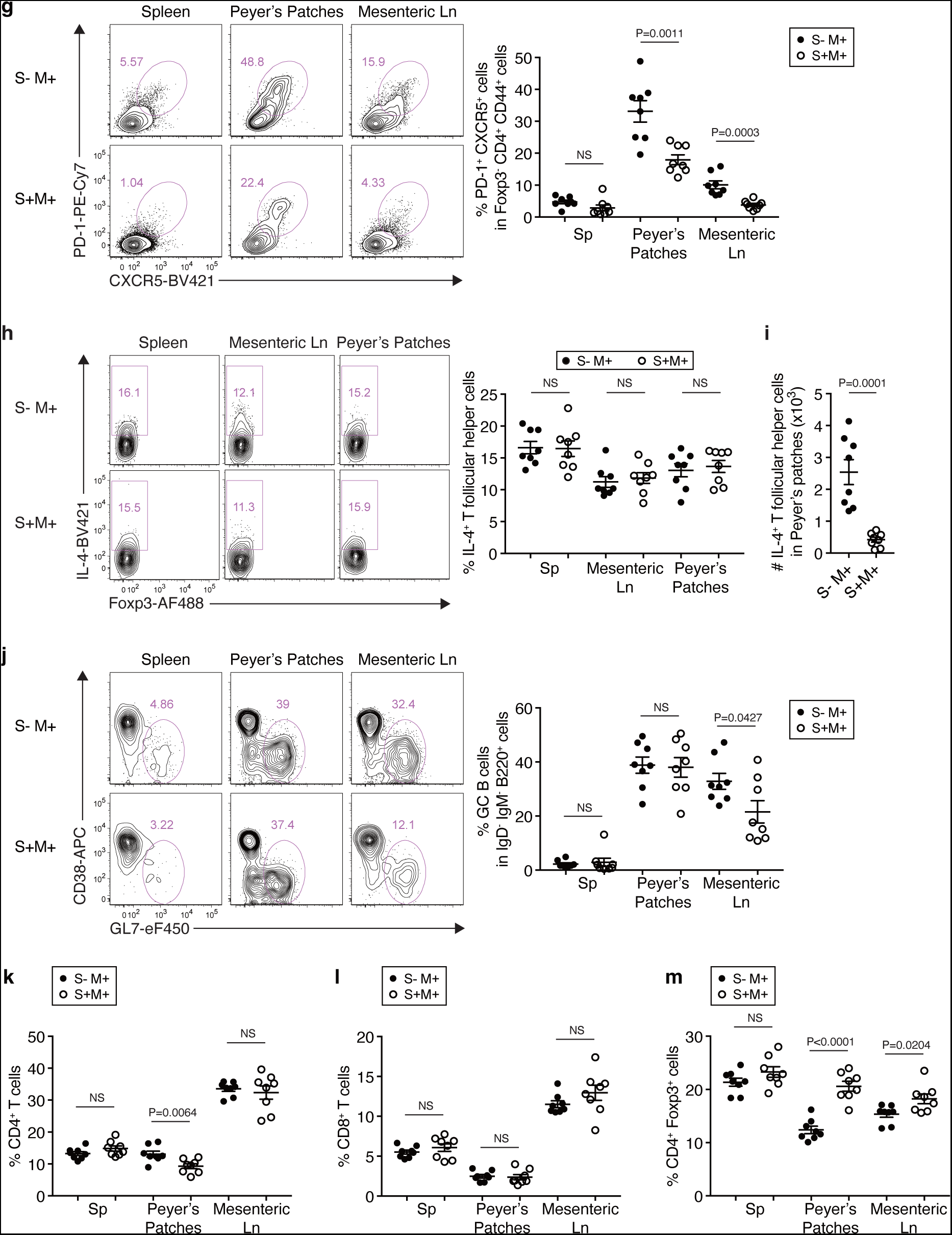
Generation of S^-^M^+^ mice and their immunological characterization. **a,** Schematic representation of a modified BAC CTLA-4 allele (RP23-146J17) in S^-^M^+^ mice. The exon3 and exon4 were conjugated to abrogate sCTLA-4 expression. **b,** Co-staining of CTLA-4 and Foxp3 in CD4^+^ T cells from 6-week-old S^-^M^+^ and S^+^M^+^ mice. **c,** Representative images of spleens and peripheral lymph nodes, and the absolute numbers of total splenocytes (Sp) and lymphocytes in mesenteric or surface lymph nodes (Ln) in 30-week-old S^-^M^+^ and S^+^M^+^ mice. **d,** HE staining of indicated organs of S^-^M^+^ and S^+^M^+^ mice. Scale bars represent 100 μm. **e,** Co-staining of CD44 and CD62L in CD8^+^ T cells from spleen (Sp) or lymph nodes (Ln) of S^-^M^+^ and S^+^M^+^ mice at 8, 16, and 24 weeks of age. Representative flow cytometry data (right panel) show the frequencies of CD44^+^ CD8^+^ T cells in 24-week-old indicated groups of mice. **f,** Co-staining of CD44 and CD62L in CD4^+^ T cells from spleens or lymph nodes of 24- week-old S^-^M^+^ and S^+^M^+^ mice. **g,** Frequencies of Tfh cells as PD-1^+^CXCR5^+^Foxp3^-^ CD44^+^CD4^+^B220^-^ cells in spleen, Peyer’s patches, and mesenteric lymph nodes of indicated groups of mice. **h,** Frequencies of IL-4-producing Tfh cells in spleen, Peyer’s patches, and mesenteric lymph nodes of S^-^M^+^ and S^+^M^+^ mice. **i,** Increase in the number of IL-4-producing Tfh cells in S^-^M^+^ mice. **j,** Frequencies of germinal center (GC) B cells as CD38^-^ GL7^+^ IgD^-^ IgM^-^ B220^+^ CD4^-^ cells in spleen, Peyer’s patches, and mesenteric lymph nodes of S^-^M^+^ and S^+^M^+^ mice. **k-m,** Frequencies of CD4^+^ T cells (**k**), CD8^+^ T cells (**l**), and CD4^+^ Foxp3^+^ cells (**m**) in spleen, Peyer’s patches, and mesenteric lymph nodes of S^-^M^+^ and S^+^M^+^ mice. Thirty-week-old mice were used for the experiments (**c,d,g-m**). Error bars denote the mean ± s.e.m. in (**c,e-m**). Data are representative or summary of at least two independent experiments with three or more mice (**b-m**). Statistical significances were determined by one-way ANOVA with Holm-Sidak multiple comparisons test in (**e, left panel**) or unpaired two-tailed *t*-test in (**c,e-m**). AF488, Alexa Fluor 488; eF450, eFluor 450.

**Extended Data Fig. 4:**
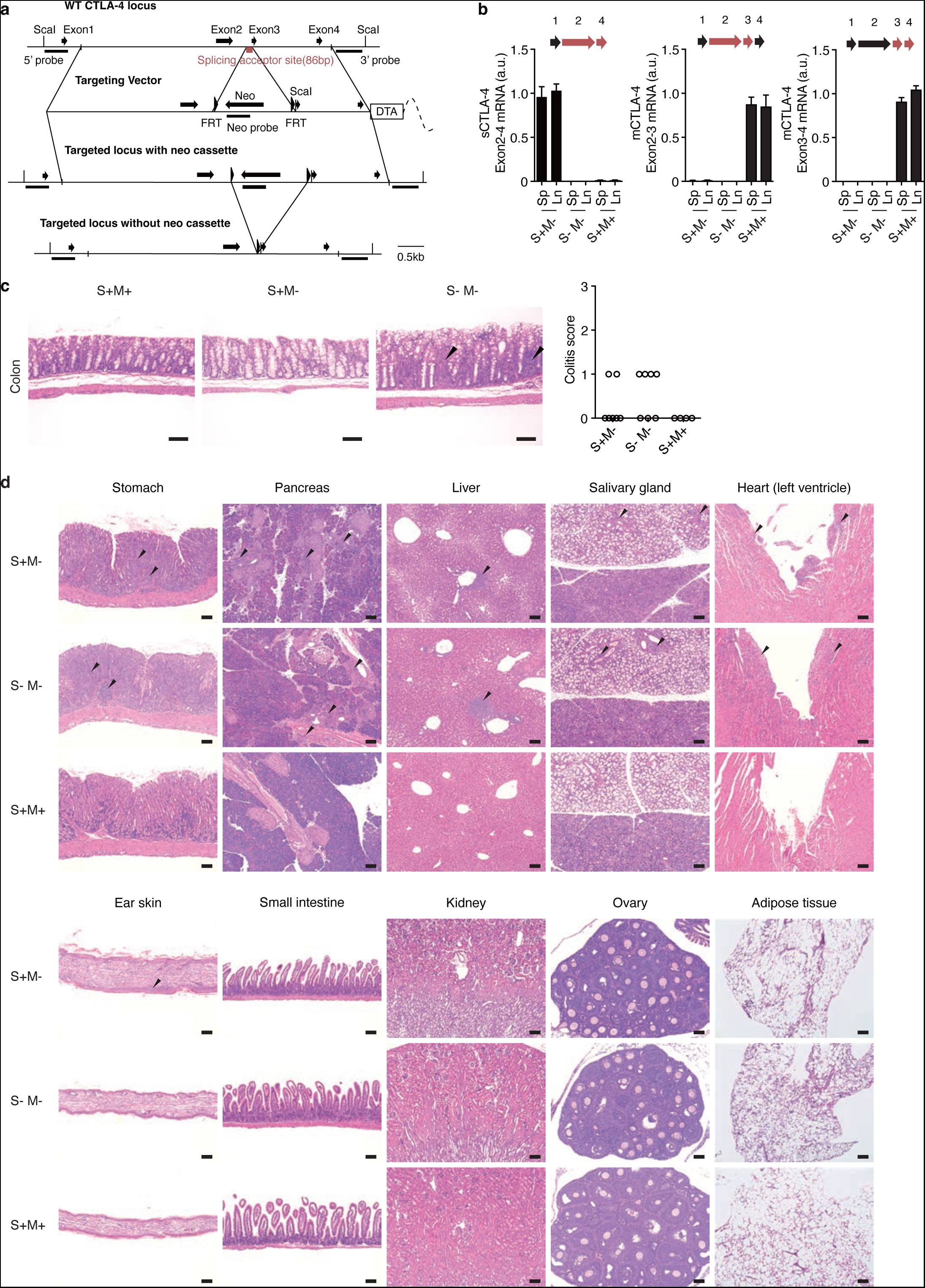
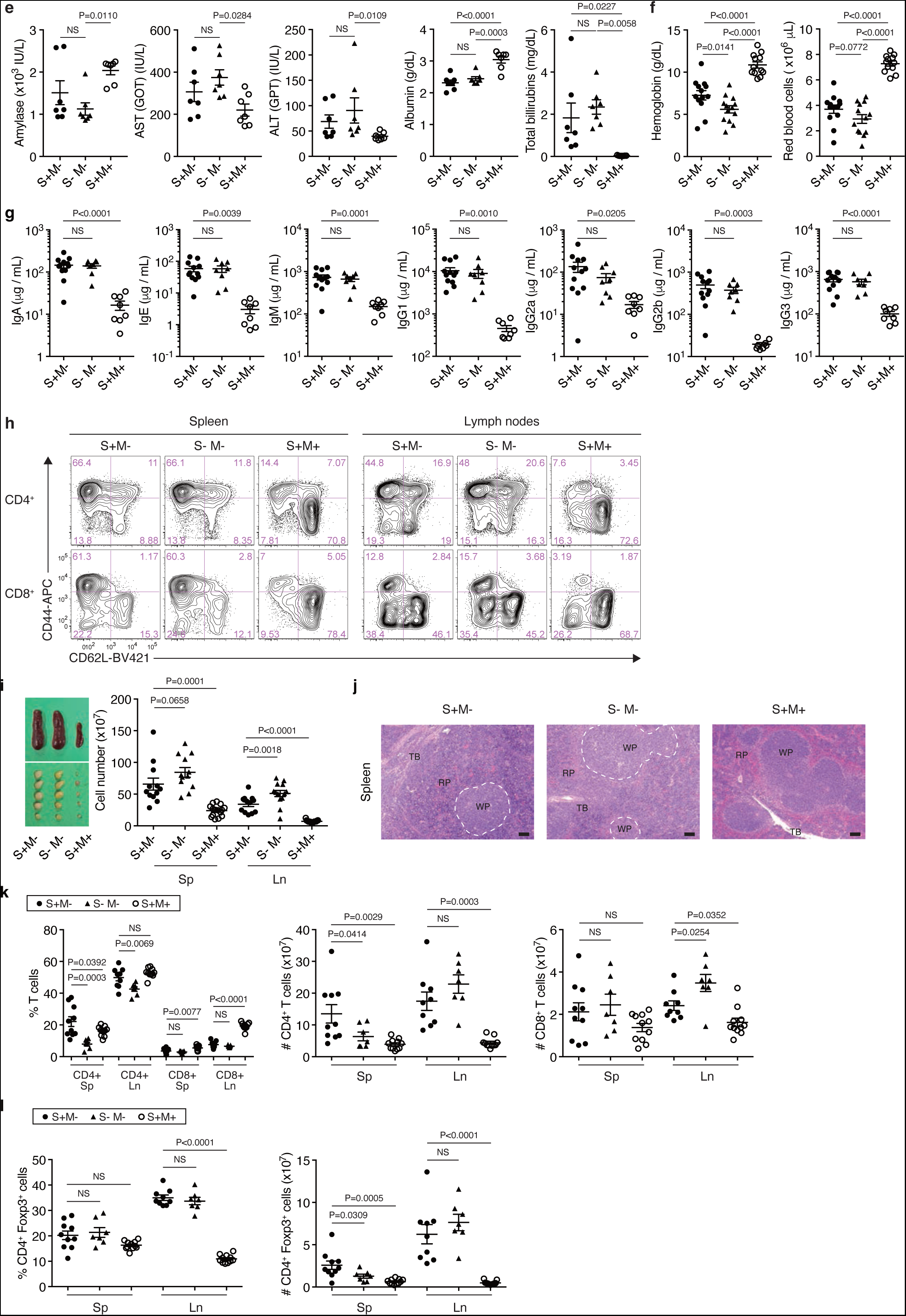
Generation of S^+^M^-^ mice and their immunological characterization. **a,** Schematic representation of the CTLA-4 allele in S^+^M^-^ mice. A targeting vector was designed to delete splicing acceptor site (86 bp) in front of the exon3. After generation of chimera mice, the mice were bred with FLPeR mice harboring FLP1 recombinase gene to delete a neo cassette. The offspring were the S^+^M^-^ mice. **b,** Constitutional exon3 skipping in CTLA- 4 mRNA in S^+^M^-^ mice. Relative mRNA expression of sCTLA-4 (exon2-4 junction spanning primers) and mCTLA-4 (exon2-3 or exon3-4 junction spanning primers) in CD4^+^ T cells of indicated groups of mice purified from spleens and lymph nodes. **c,** HE staining of colons of indicated groups of mice (n = 4 to 7 per group). Scale bars represent 100 μm. Colitis were scored histologically. The data were assessed by Kruskal-Wallis test followed by Dunn’s test. **d,** HE staining of indicated organs of S^+^M^-^, S^-^M^-^, and S^+^M^+^ mice (n = 6 per group). Scale bars represent 100 μm. Arrows show leukocyte infiltration in tissues. **e,** Blood chemistry analysis for assessment of organ functions and inflammation. The data of ALT was accessed by Kruskal-Wallis test followed by Dunn’s test. AST, aspartate aminotransferase; ALT, alanine transaminase. **f,** Decrease in serum hemoglobin concentration and red blood cell counts in S^+^M^-^ and S^-^M^-^ mice. **g,** Serum concentrations (μg/mL) of IgA, IgE, IgM, IgG1, IgG2a, IgG2b, and IgG3 immunoglobulins in indicated groups of mice. **h,** Representative data of CD44/CD62L expression by CD4^+^ or CD8^+^ T cells in S^+^M^-^, S^-^M^-^, and S^+^M^+^ mice shown in Fig. 3j. **i,** Splenomegaly and lymphadenopathy in S^+^M^-^ and S^-^M^-^ mice. **j,** HE staining of spleens of S^+^M^-^, S^-^M^-^, and S^+^M^+^ mice (n = 6 per group). White pulps in S^+^M^-^ and S^-^M^-^ mice are shown by white dot lines because of obscure borders by massive leukocyte infiltration into red pulps. WP, white pulp; RP, red pulp; TB, trabecular vessel. Scale bars represent 100 μm. **k,** Ratios and numbers of CD4^+^ and CD8^+^ T cells in spleens/lymph nodes of indicated groups of mice. **l,** Ratios and numbers of Foxp3^+^CD4^+^ T cells in spleens/lymph nodes of indicated groups of mice. Three-week-old mice were used for the experiments (**b-l**) because of the early mortality of S^-^M^-^ mice. Error bars denote the mean ± s.e.m. in (**e-g,i,k,l**) or the mean ± s.d. in (**b**). Data are representative or summary of at least two independent experiments with three or more mice (**b-l**). Statistical significances were determined by one-way ANOVA with Holm-Sidak multiple comparisons test in (**e-g,i,k,l**). Neo, Neomycin resistance cassette; FRT, Flippase Recognition Target.

**Extended Data Fig. 5:**
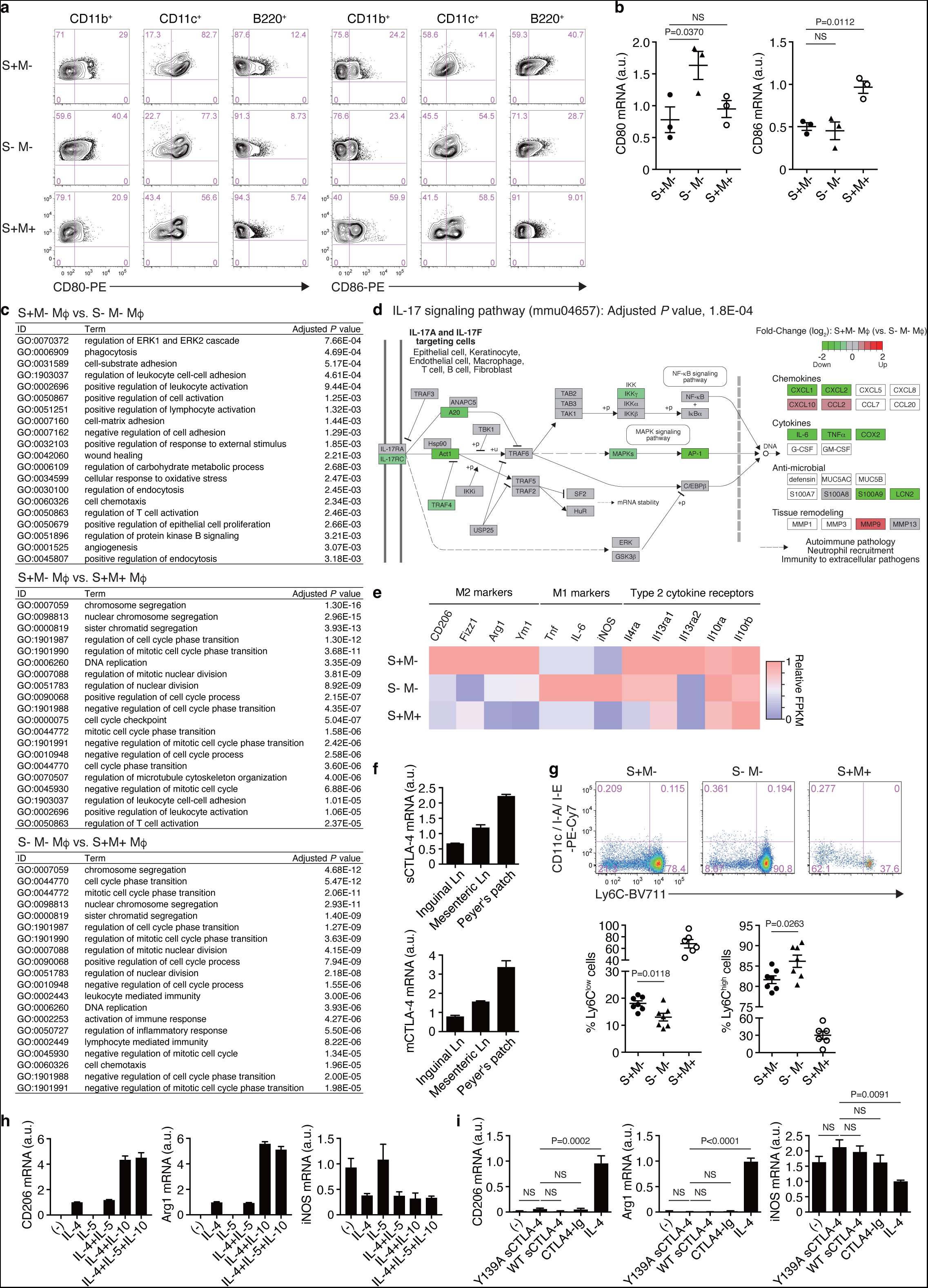
sCTLA-4 induces M2-like macrophage differentiation under chronic inflammation. **a,** CD80 and CD86 expression by CD11b^+^, CD11c^+^, and B220^+^ cells in spleen of indicated groups of mice. Data were representative of flow cytometric analysis shown in Fig. 4a (n = 8 to 14 per group). **b,** Relative mRNA expression of CD80 and CD86 in CD11b^+^ splenocytes from indicated groups of mice. **c,** Gene ontology (GO) enrichment analysis on biological process. The top sets of enriched GO terms (over-represented) in comparisons between indicated macrophage (Mϕ) groups were shown. To assess statistical significances, Benjamini-Hochberg method was applied to obtain adjusted *P* values for multiple testing. **d,** Top enriched KEGG pathway in differentially expressed genes (DEGs) downregulated in S^+^M^-^ peritoneal macrophages compared with S^-^M^-^ ones (mmu04657, IL-17 signaling pathway). Green squares indicate differentially down-regulated in S^+^M^-^ macrophages (Log with base 2 fold-change). **e,** Relative FPKM values of M1/M2-related genes and type 2 cytokine receptors in F4/80^+^CD11b^+^ peritoneal macrophages purified from indicated groups of mice. The values are illustrated by color-coded matrix. **f,** Relative expression of sCTLA-4 and mCTLA-4 in CD4^+^ T cells from indicated lymph nodes. **g,** Frequencies of Ly6C^lo^ and Ly6C^hi^ monocytes as CD45^+^ CD11b^+^ CD11c^-^ F4/80^-^ I-A/I-E^-^ Ly6G^-^ Siglec-F^-^ CD90^-^ B220^-^ CD49b^-^ blood cells prepared from S^+^M^-^ and S^-^M^-^ mice. Unpaired two-tailed *t*-test was applied to analyze the difference in blood monocytes between S^+^M^-^ and S^-^M^-^ mice. **h,i,** Relative mRNA expression for CD206, Arg1, and iNOS in RAW264.7 cells 24 hours after treatment of IL-4 (10 ng/mL), IL-5 (10 ng/mL), IL-10 (10 ng/mL), WT sCTLA-4 (5 μg/mL), Y139A sCTLA-4 (5 μg/mL), or CTLA4-Ig (5 μg/mL). Independent experiments were performed two (**h**) or three times (**i**). Three-week-old mice were used for the experiments (**a-e,g**). Error bars denote the mean ± s.e.m. in (**b,g-i**) or the mean ± s.d. in (**f**). Data are representative or summary of at least two independent experiments with two or more mice per group (**a,b,f,g**), or RNA-seq experiments with two mice per group (**c-e**). Statistical significances were determined by one-way ANOVA with Holm-Sidak multiple comparisons test in (**b,i**). BV711, Brilliant Violet 711.

**Extended Data Fig. 6:**
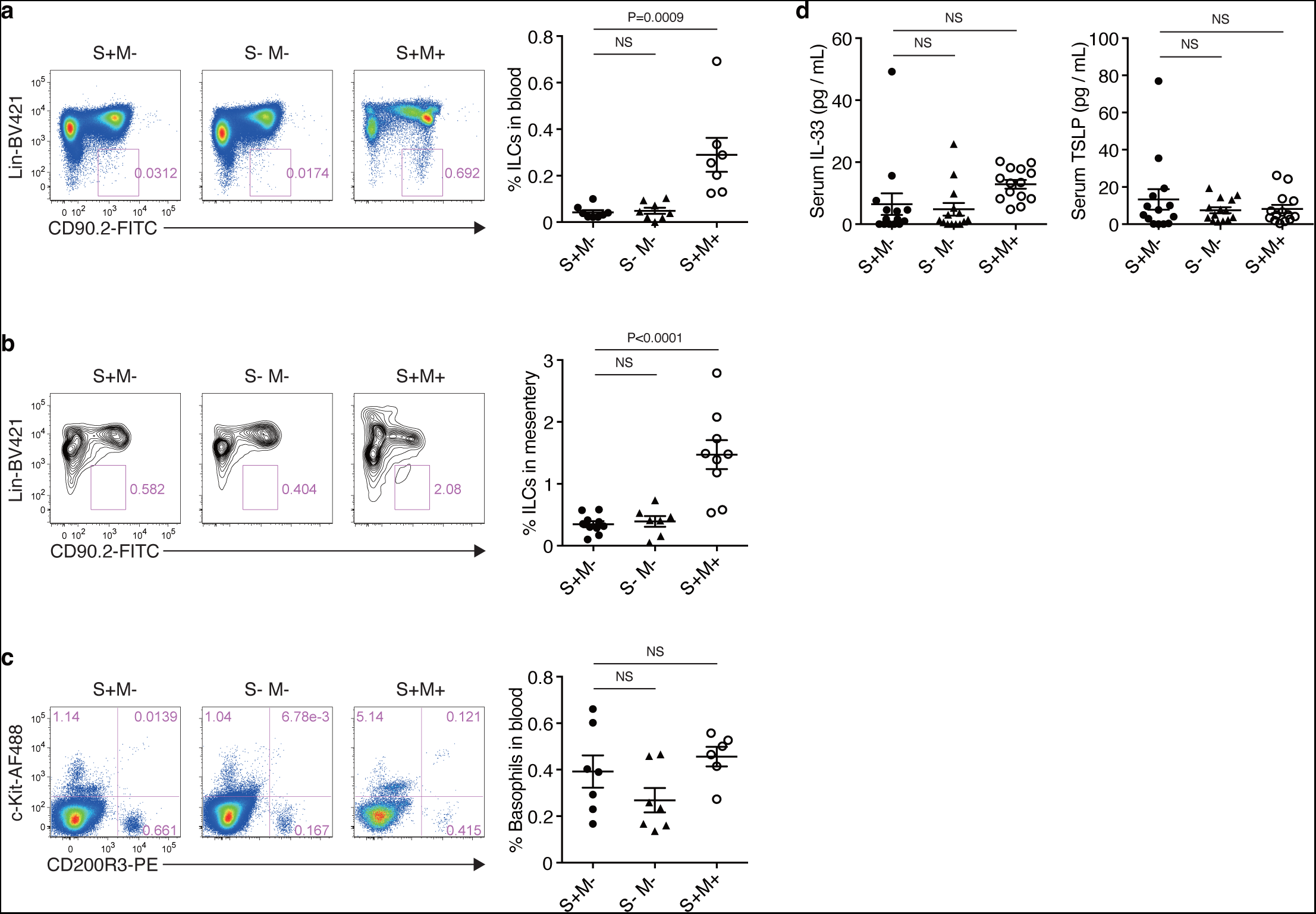
Non-increase in serum concentration of IL-33 and TSLP, and in the proportions of ILCs, basophils, and mast cells. **a,b,** Frequency of ILCs as CD45^+^Lin^-^ (Lineage: CD3, CD4, CD8a, CD11b, CD11c, CD19, CD49b, Gr-1, TER119, FcεRIα, F4/80) CD90.2^+^ cells in blood (**a**) and mesentery (**b**) prepared from indicated groups of mice. **c,** Frequency of c-Kit^+^ cells and basophils as c-Kit^-^CD200R3^+^ cells in CD45^int^ blood cells prepared from indicated groups of mice. **d,** Serum concentrations of IL-33 and TSLP in indicated groups of mice. Three-week-old mice were used for all the experiments (**a-d**). Error bars denote the mean ± s.e.m. in (**a-d**). Data are representative or summary of at least two independent experiments with six or more mice (**a-d**). Statistical significances were determined by one-way ANOVA with Holm-Sidak multiple comparisons test in (**a-d**). ILCs, innate lymphoid cells.

**Extended Data Fig. 7:**
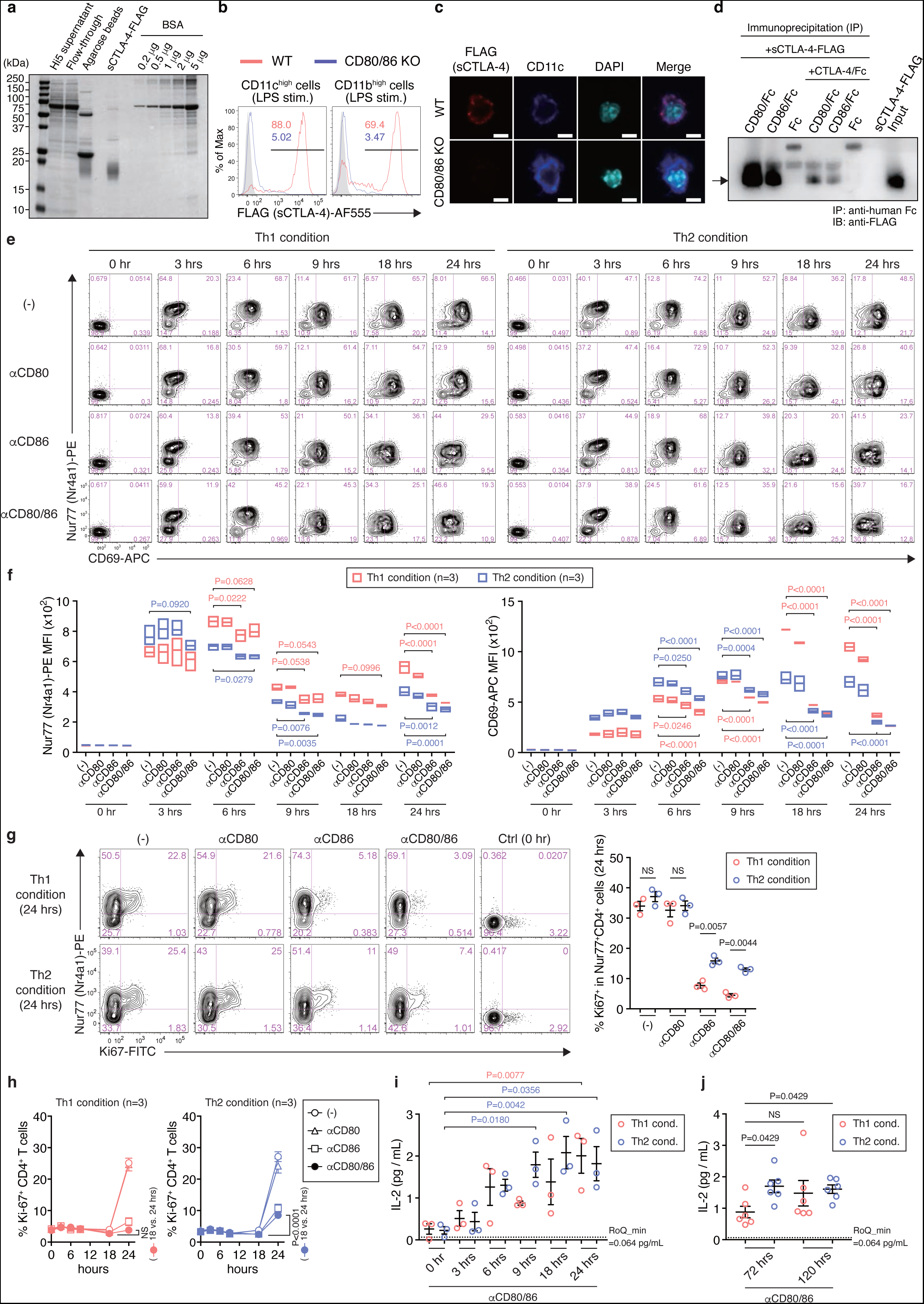
CD80/86-binding capacity of recombinant sCTLA-4 protein and its effects on *in vitro* Th1/Th2 differentiation. **a,** sCTLA-4-FLAG recombinant protein. Purified protein was detected by SDS-PAGE and CBB staining. Baculovirus expression system was used for expression of the recombinant protein. **b,** Staining of FLAG in LPS-stimulated CD11b^hi^ or CD11c^hi^ cells pre-incubated with sCTLA-4-FLAG (1 μg/mL, for 20 minutes at 4°C). The CD11b^hi^ and CD11c^hi^ cells were prepared from WT or CD80/86 KO mice. **c,** Immunofluorescence microscopy of splenic CD11c^+^ cells with or without CD80/86 expression pre-incubated with sCTLA-4-FLAG and then stained for FLAG (Alexa fluor 555), CD11c (Alexa fluor 647), and DAPI. Scale bar, 5 μm. **d,** Weaker affinity of sCTLA-4 for CD80/86 compared with CTLA-4 Fc chimera. 1 μg FLAG-tagged sCTLA-4 and 1 μg CD80 or CD86 Fc chimera protein (Fc: human IgG1) were immunoprecipitated with anti-human Fc antibody-coated Dynabeads in the presence or absence of 1 μg CTLA-4 Fc chimera (Fc: mouse IgG2a). Recombinant Fc (Fc: human IgG1) was used as a negative control. The immunoprecipitates were analyzed by immunoblotting with anti-FLAG M2 antibody. Data are representative of two independent experiments (**a-d**). **e,f,** Comparison of Nur77 (Nr4a1) and CD69 expression kinetics in Th1 and Th2 cells skewed in the presence of αCD80/86 antibodies. **g,** Comparison of Ki-67 expression in Nur77 positive cells in Th1 and Th2 differentiation 24 hours after TCR stimulation. **h,** Ki-67 expression kinetics under Th1 and Th2 differentiation during 0-24 hours after TCR stimulation. Each symbol indicates the presence or absence of αCD80 or αCD86 antibody, or both. **i,j,** Quantification of IL-2 by PEA-qPCR (lower limit of quantification, 0.064 pg/mL, as shown in the dotted line) in the presence of αCD80/CD86 antibodies for 0-24 hours in Th1/Th2 polarization experiments shown in **e-h** (**i**), and quantification of IL-2 levels at 72 and 120 hours after TCR stimulation (**j**). Two-way ANOVA with Holm-Sidak multiple comparisons test (**f-i**). Kruskal-Wallis test with Dunn’s multiple comparisons test (**j**). Data points show results of three independent experiments with cells isolated from different mice (n = 3 to 6 mice per experiment) (**f-j**). PEA, Proximity Extension Assay.

**Extended Data Fig. 8:**
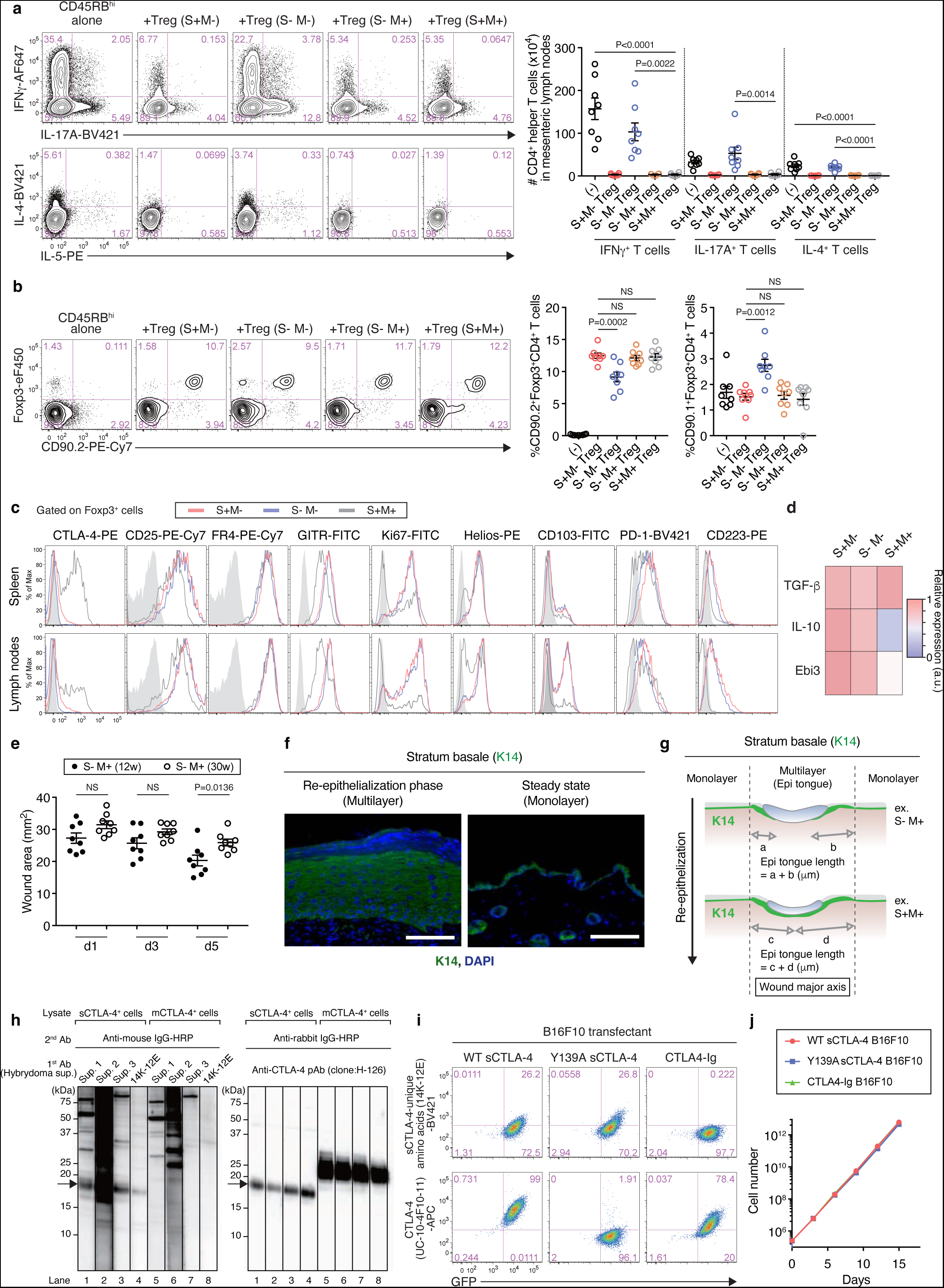
*In vivo* immuno-suppressive and -modulatory effects of sCTLA-4 on intestinal inflammation, tumor immunity, and tissue repair. **a,** IFNγ^+^ T cells (Th1), IL-17A^+^ T cells (Th17), and IL-4^+^ T cells (Th2) in draining mesenteric lymph nodes 8 weeks after adoptive cell transfer as shown in Fig. 7a. Representative staining of CD4^+^ T cells for cytokines and the absolute numbers of Th1, Th17, and Th2 cells are shown. **b,** Foxp3^+^ Tregs in mesenteric lymph nodes 8 weeks after adoptive transfer as shown in Fig. 7a. Representative Foxp3 staining and the frequencies of Foxp3^+^CD4^+^ T cells derived from transferred CD90.1^+^(Thy1.1^+^)CD45RB^hi^ Tconvs and those derived from CD90.2^+^(Thy1.2^+^) Tregs co-transferred with the Tconvs are shown. **c,** Expression of Treg-associated and activation-associated molecules in Foxp3^+^CD4^+^ T cells from indicated groups of mice. **d,** Relative expression of mRNA for TGF-β, IL-10, and Ebi3 determined by qPCR in Foxp3^+^(GFP^+^)CD4^+^ T cells purified from DEREG S^+^M^-^, DEREG S^-^M^-^, and DEREG S^+^M^+^ mice (n = 3 per group). **e,** Comparison between 12-week-old S^-^M^+^ mice and 30-week-old S^-^M^+^ mice in the degree of wound closure. Average of duplicate wound area (mm2) per mouse was measured as described in Fig. 7c,d. Scale bars represent 2 mm. **f,g,** Quantification of re-epithelialization by the length of the tongue (Epi tongue length). K14^+^ epidermal keratinocytes in the wound sites proliferate, become multilayered, and migrate into under the wounds from the excision edges (**f**). The edge of the wounds on day 5 was defined as the boundary between K14^+^ monolayer and multi-layer keratinocytes. An example of the measurement is shown in (**g**). Scale bars represent 100 μm. **h,** sCTLA-4-specific mAb. Specificities to sCTLA-4 of 4 supernatants (sup.) from cloned hybrydomas after screening were confirmed by the Western blots using sCTLA-4^+^ or mCTLA-4^+^ transfectants. A mAb (clone: 14K-12E) specifically detected sCTLA-4 but not mCTLA-4 (left panel). The mAbs as the culture supernatants were stripped off from the Western blotting membranes; sCTLA-4/mCTLA-4 were re-probed by the CTLA-4 polyclonal antibody (H-126) to assess each exact size (right panel). The lanes were fragmented because each lane was separately stained by the different hybridoma supernatants. The antibody (14K-12E) was only effective on sCTLA-4 over-expressing cell lines. **i,** B16F10 melanoma cells secreting WT sCTLA-4, Y139A sCTLA-4, or CTLA4-Ig. The cells were generated by retroviral transduction using pMCsIg-WT sCTLA-4, pMCsIg-Y139A sCTLA-4, and pMCsIg-CTLA4-Ig. Infected cells were sorted as GFP^+^ cells to prepare transduced cells. To confirm protein production by the transfectants, the transduced cells were stained with anti-CTLA-4 mAb (UC10-4F10-11) or mAb specific for sCTLA-4-unique amino acids (14K-12E), and analyzed by flow cytometry. The 14K-12E mAb did not recognize CTLA4-Ig; the UC-10- 4F10-11 mAb did not react with the sCTLA-4 protein with Y139A mutation in MYPPPY motif. **j,** *In vitro* growth of B16F10 melanoma transfectants. Growth curves were determined after at least five times of consecutive cell passages *in vitro*. Three-week-old mice were used for the experiments (**c,d**). Error bars denote the mean ± s.e.m. in (**a,b,e**). Data are representative or summary of at least two independent experiments with two or more mice per group (**a-f**). Statistical significances were determined by one-way ANOVA with Holm-Sidak multiple comparisons test in (**a,b**) or unpaired two-tailed *t*-test in (**e**).

**Extended Data Fig. 9:**
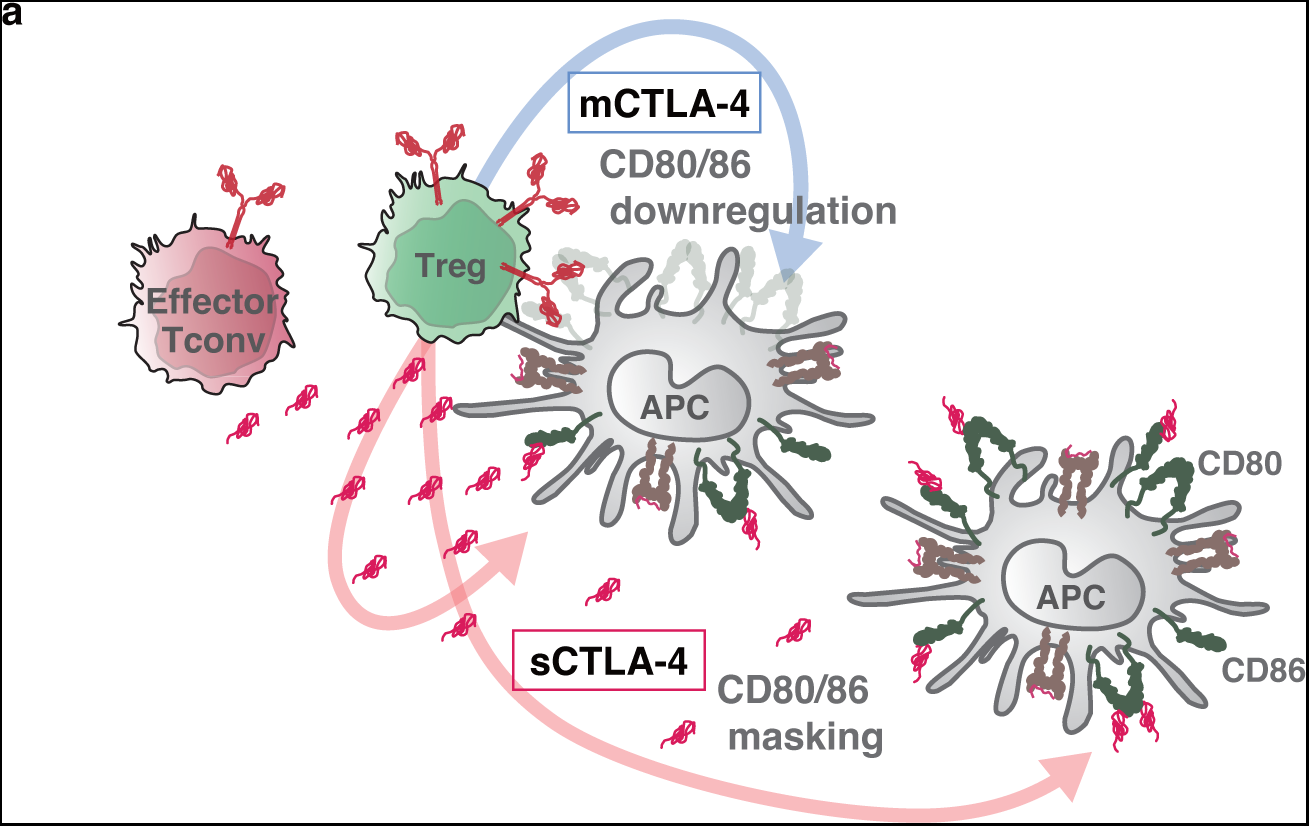
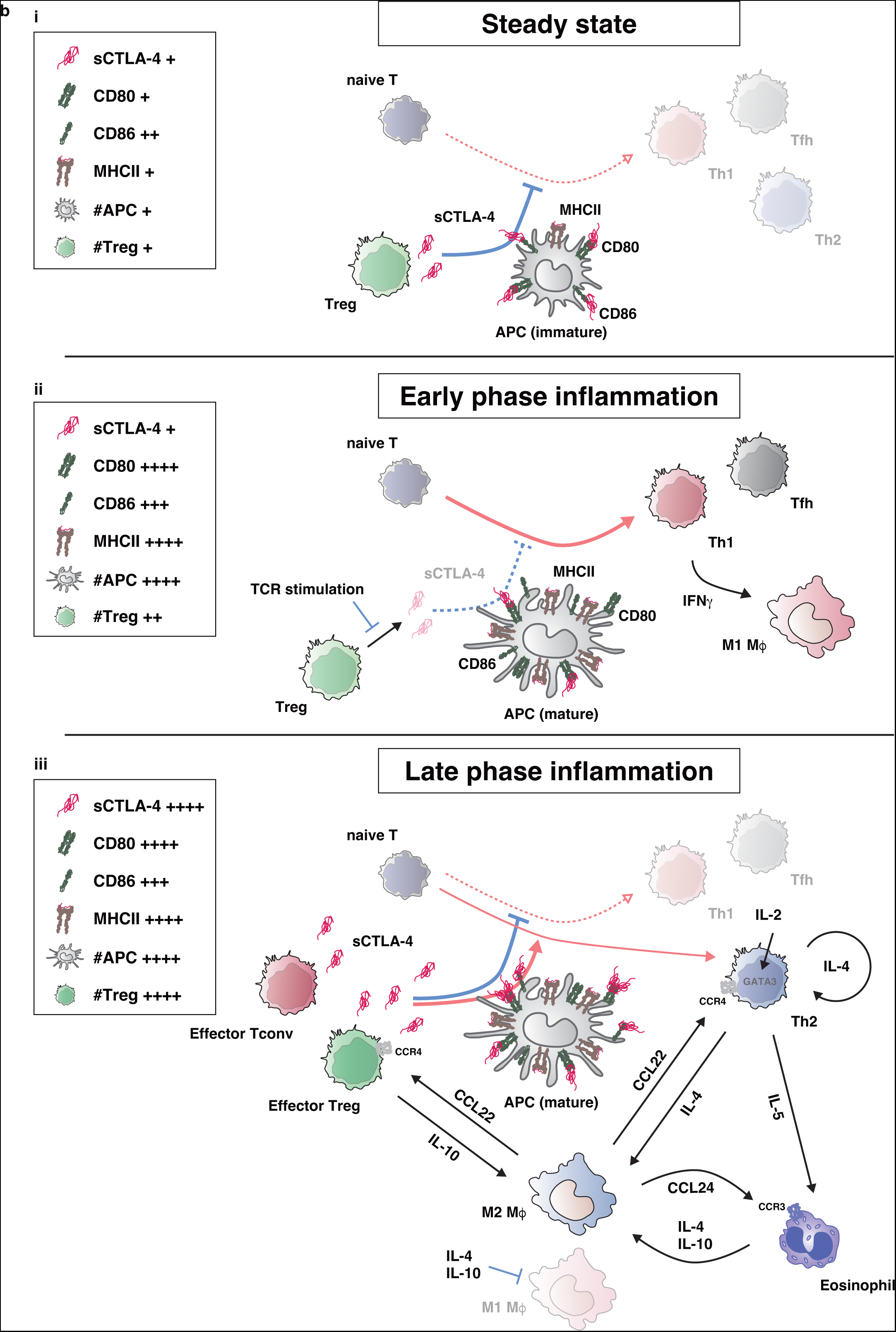
The roles of sCTLA-4 for systemic and local immunoregulation in steady and inflammatory states. **a,** The roles of sCTLA-4 in immune suppression. In Treg-conjugated APCs, Treg mCTLA-4 down-modulates CD80/CD86 expression, and sCTLA-4 produced by Tregs and nearby activated Tconvs blocks residual CD80/CD86 to augment suppression. sCTLA-4 also blocks CD80/CD86 expressed on Treg-nonconjugated APCs in the vicinity, exerting bystander suppression. **b,** Contribution of sCTLA-4 to immune homeostasis in steady and inflammatory states. **(i) Steady state.** In order to prevent the activation of self-reactive Tconvs in the steady state, sCTLA-4 mainly produced by Tregs is raising the T-cell activation threshold locally and systemically by masking CD80/86 molecules expressed on immature antigen presenting cells (APCs) that are CD80/CD86^lo^ and MHCII^lo^. **(ii) Early phase inflammation.** In early stages of inflammation when activated Tregs with high mCTLA-4 expression are not much recruited to the inflammation site, temporary down-regulation of sCTLA-4 production by TCR-stimulated Tregs facilitates naive T cells to be primed by activated APCs that are CD80/CD86^hi^ and MHCII^hi^, and differentiate into Th1/Th17/Tfh cells mediating immune responses. **(iii) Late phase inflammation.** In active and late stages of inflammation, effector Tregs as well as effector Tconvs are recruited to the inflammation site, become activated to up-regulate sCTLA- 4 production as well as mCTLA-4 expression. Abundantly produced sCTLA-4 blocking co-stimulation suppresses the differentiation of Tconvs towards Th1/Tfh cells and allows the differentiation towards Th2 cells, which enhance M2-like macrophage differentiation along with eosinophil activation via Th2 cytokines (such as IL-4 and IL-10) to prevent excessive host damage, resolve inflammation, and promote tissue repair and remodeling.

**Extended Data Fig. 10:**
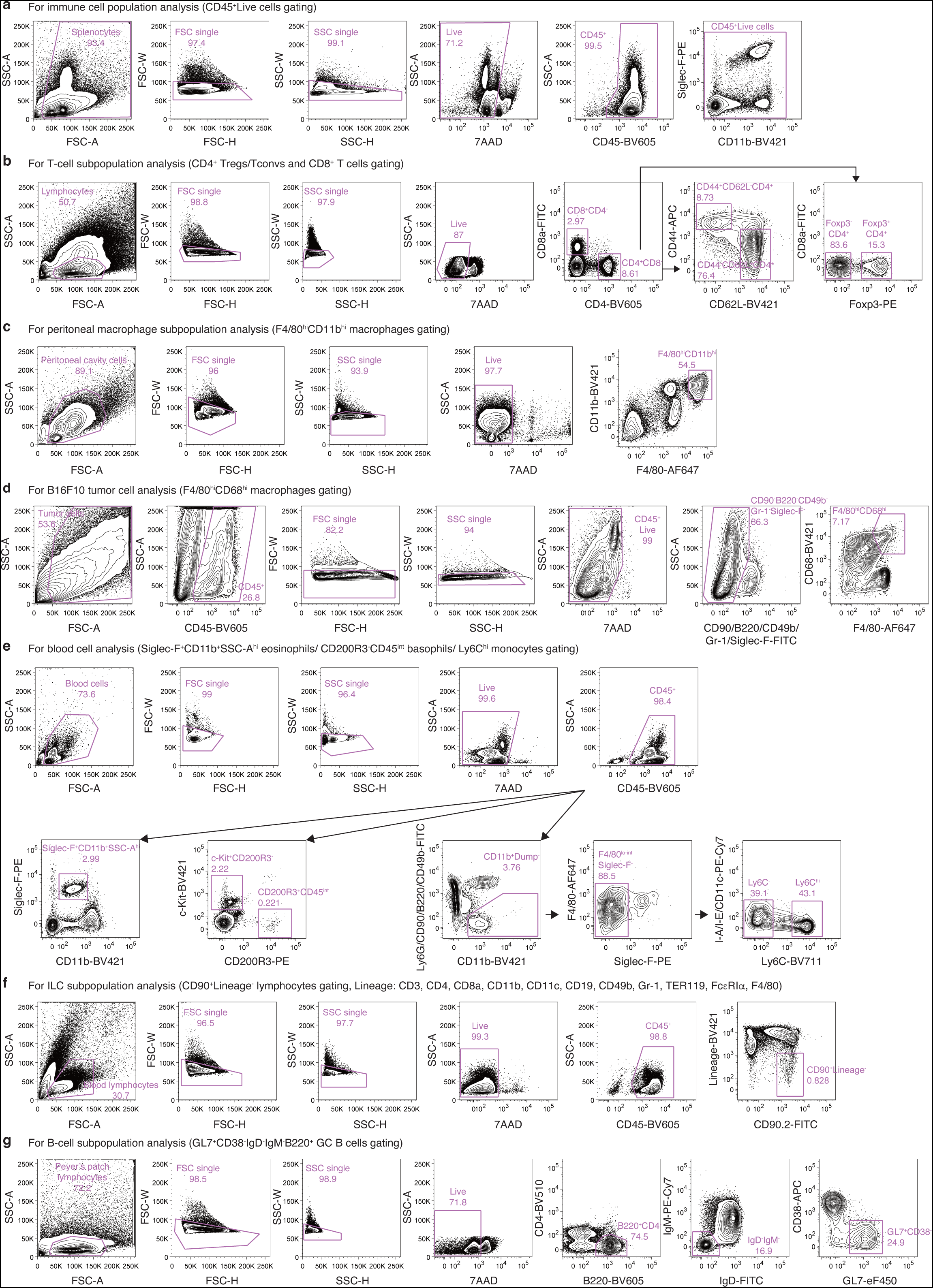
Gating strategies for murine immune cell studies. **a-g,** Representative flow cytometry data show gating strategies to obtain single CD45^+^ live cells for immune cell population analysis (**a**), Foxp3^+/-^, CD44^+/-^, or CD62L^+/-^ CD4^+^ T cells and CD8^+^ T cells for T-cell subpopulation analysis (**b**), F4/80^hi^CD11b^hi^ macrophages for peritoneal macrophage subpopulation analysis (**c**), B16F10-infiltrating single CD45^+^ live cells for immune cell population/subpopulation analysis (e.g. F4/80^hi^CD68^hi^CD11b^+^ macrophages) (**d**), eosinophils, basophils, mast cells, or Ly6C^+/-^ monocytes for blood cell analysis (**e**), CD90^+^Lineage^-^ lymphocytes for ILC subpopulation analysis (**f**), and germinal center B cells for B-cell subpopulation analysis (**g**). BV510/605, Brilliant Violet 510/605.

